# Function-specific epistasis shapes evolutionary trajectories towards antibiotic resistance

**DOI:** 10.1101/2025.07.09.663857

**Authors:** Gabriela Petrungaro, Theresa Fink, Booshini Fernando, Gerrit Ansmann, Tobias Bollenbach

## Abstract

Pre-existing mutations can influence subsequent evolution by constraining or opening evolutionary pathways through epistatic interactions. In some cases, global epistasis allows evolutionary pathways to be predicted from the fitness of the genetic background alone. In other cases, idiosyncratic epistasis makes evolution less predictable. Here, we show that the evolution of antibiotic resistance is highly repeatable, following a common path across most genetic backgrounds. However, a minority of genetic backgrounds exhibit evolutionary trajectories that significantly deviate from this common path. Rather than being predictable from global epistasis, these deviations are modulated by function-specific epistasis: perturbations to specific cellular functions lead to novel evolutionary trajectories towards resistance. Using tightly controlled robotic evolution experiments, we quantitatively analyzed resistance trajectories for three clinically relevant antibiotics across multiple genetic backgrounds, including hundreds of *Escherichia coli* gene-deletion strains and several clinical isolates from urinary-tract infections. We show that disrupting distinct sets of cellular functions alters evolutionary trajectories for specific drugs and across different drugs, and we identify genetic changes defining these alternative trajectories. Importantly, this function-specific epistasis often slows down resistance evolution. Some of these effects can also be induced by small-molecule inhibitors of the identified targets, suggesting that function-specific epistasis can be exploited to improve drug treatments.

## Introduction

The ubiquitous influence of epistatic interactions between mutations intricately shapes the course of microbial evolution. In extreme cases, the presence of a single mutation can completely alter the fitness effects of subsequent mutations and the resulting mutational path^1–3^, thereby constraining or potentiating future evolution^4,5^. Such idiosyncratic epistasis generally reduces the predictability of evolution, but in principle makes it possible to redirect evolution through targeted intervention (Fig. S1a). Alternatively, global patterns of epistasis – such as diminishing-returns, where the effects of beneficial mutations decrease in fitter genetic backgrounds – may predominate^6–11^ (Fig. S1b). Global epistasis patterns can result from inherent nonlinearities in fitness landscapes^12–14^. Evolutionary outcomes are then largely determined by initial phenotypes, resulting in convergent phenotypic evolution^8,15,11,16^. This would facilitate evolutionary prediction, but also implies that fundamental changes in evolutionary trajectories are hard to achieve.

These phenomena are particularly relevant in the context of antibiotic resistance. In addition to being a serious public-health problem^17^, resistance evolution serves as an excellent model system to study fundamental evolutionary questions^7,18–31^. This is due to the relatively short timescales and readily quantifiable phenotypes: spontaneous mutations and antibiotic selection can increase resistance by orders of magnitude within a week^31,18^. Recent work has revealed contrasting patterns of epistasis in this context. For example, the evolution of resistance to a ribosome-inhibiting antibiotic exhibits a global epistasis pattern, where the effect of gene deletions on initial resistance largely predicts subsequent resistance gains^26^. In contrast, a study using clones from the *E*. *coli* Long-Term Evolution Experiment has identified idiosyncratic epistasis in antibiotic resistance^19^. While there are a few known cases where pre-existing mutations alter evolutionary trajectories to resistance^2,7,26,32^, the prevalence of this phenomenon, the specific cellular functions involved, the role of global epistasis versus idiosyncratic epistasis, and the potential to exploit such effects to slow down resistance evolution are largely unknown.

The search for gene deletions that can alter, and in particular suppress or slow down, the evolution of resistance is an active area of research^21,26,33^. In *P. aeruginosa,* deleting *ampC* and *ampR* dramatically reduces the ability to evolve resistance to the antibiotic ceftazidime^21^. Other work found that the global regulator *ampR* and the quorum-sensing regulator *lasR* strongly affect the ability to evolve resistance to specific antibiotics^3^. However, due to the differences between bacterial species and the stochastic nature of the evolutionary process, quantitative studies of epistasis and contingency in resistance evolution require tightly controlled evolution experiments with precise measurements of phenotypic adaptation over time in different genetic backgrounds and multiple replicates. Laboratory automation of evolution experiments has made it possible to scale up replicate populations, enabling more systematic studies involving hundreds of gene-deletion strains^26,33^. Tight control and precise measurements have been achieved with the *morbidostat*, a custom-built device that controls population size and selection pressure in resistance evolution experiments^31^. This technique focuses on spontaneous mutations, which, along with horizontal gene transfer, are a major mode of evolution of clinically relevant antibiotic resistance^17,34^. Recently scaled up for higher throughput using robotics, this technique has enabled the discovery that specific genes, such as the chaperone *dnaK*, alter *E. coli’*s ability to evolve resistance to ribosome-targeting antibiotics^26^. This result spurred the discovery of a small-molecule inhibitor of DnaK^35^, which in principle could serve as an “anti-evolution” drug. However, a systematic study of the effects of different genotypes on resistance evolution to clinically relevant antibiotics with different modes of action is currently lacking, partly due to significant technical challenges. As a result, it remains unclear whether perturbations of specific functions affect the ability to evolve resistance broadly or if their effects are specific to each antibiotic.

To systematically characterize the effects of genotype on phenotypic resistance evolution and mutational paths, we evolved hundreds of *E. coli* populations with different genotypes under clinically relevant antibiotics with distinct modes of action. Despite the stochastic nature of evolution, overall resistance evolution was highly repeatable at both mutational and phenotypic levels, even across different genetic backgrounds. We found that the remaining variability in resistance gain was mainly due to differences in genetic background, but not explained by global epistasis. Notably, disruption of specific cellular functions dramatically suppressed resistance evolution, even after weeks under high selection pressure. In extreme cases, this function-specific epistasis (Fig. S1c) diverted evolution away from the usual mutational paths, forcing it to explore uncharted territory. Approved drugs that target the identified functions modestly affect resistance increases in evolution experiments where they are employed together with antibiotics, showing that function-specific epistasis could be exploited to slow resistance evolution in applications.

## Results

### High-throughput platform enables quantitative comparison of nearly a thousand parallel well-controlled evolution experiments

To compare many evolution experiments under tightly controlled conditions, we scaled up a high-throughput version of the morbidostat^26,31^ (Methods). Briefly, we monitor the bacterial population size through optical density (OD) and control population size by diluting the cultures to a fixed OD every 4.2 hours. High selection pressure for antibiotic resistance is dynamically maintained by a feedback control loop that adjusts the drug concentration at each dilution to achieve 50% growth inhibition relative to the ancestor in drug-free growth medium, i.e., the IC₅₀. We implemented this technique using a dedicated robotic platform (Methods; Fig. 1a). Bacterial cultures grow in 96-well plates at 30°C and are kept in the exponential growth phase throughout the entire experiment (Fig. 1a,b). The concentration in each well is a precise measure of the resistance (IC₅₀) of the population (Fig. 1c,d; Fig. S2)^26^. Our platform can run 864 parallel evolution experiments (corresponding to nine 96-well plates) for weeks at a time, achieving approximately 160 generations per week. Contamination, a notorious problem in evolution experiments, is almost completely eliminated by several preventive measures (Methods). Compared to previous work^26^, we have more than doubled both the throughput and the maximum run time, enabling the systematic evaluation of repeatability and general patterns in evolution.

**Fig. 1:**
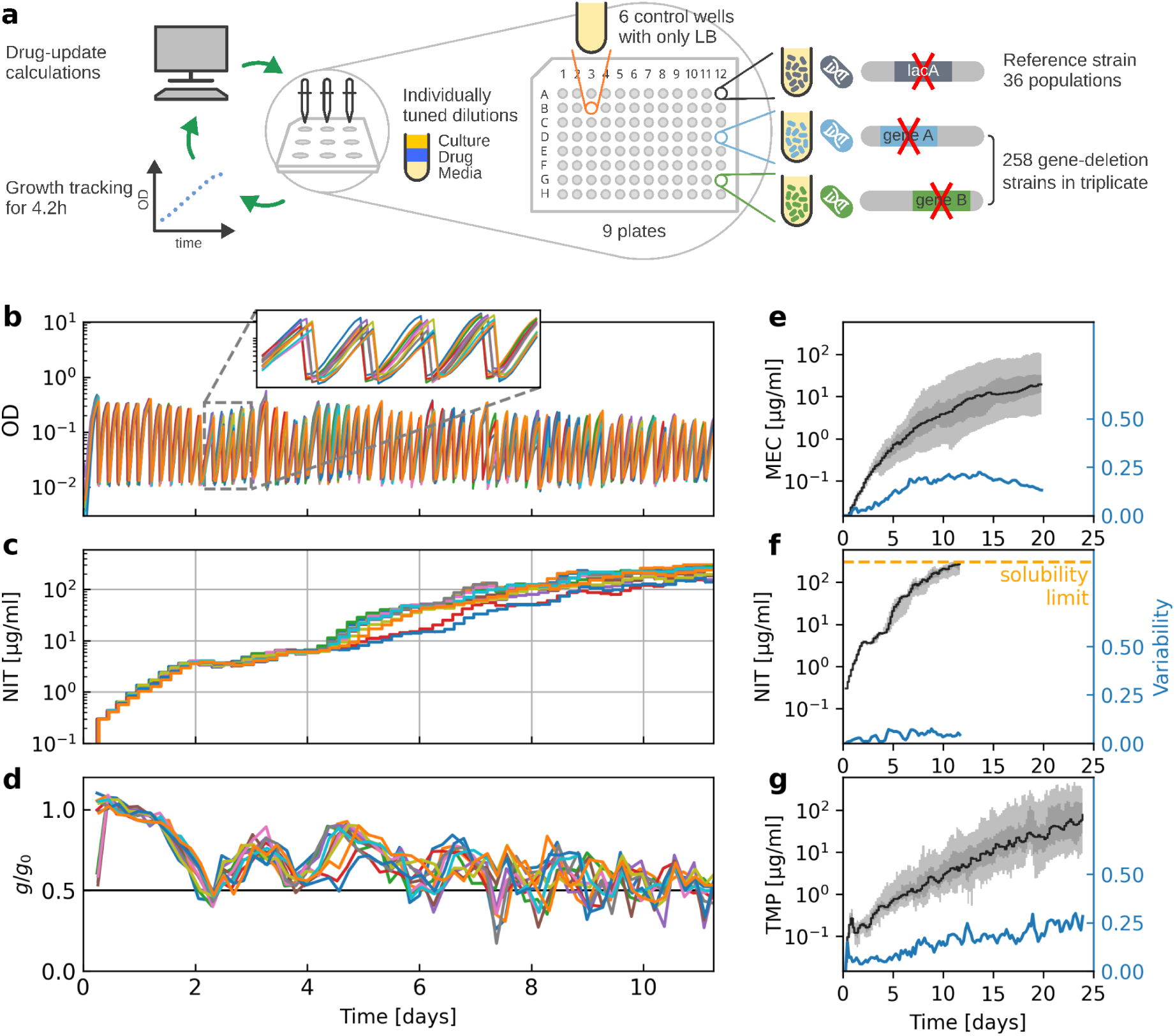
Phenotypic trajectories of resistance evolution to clinically relevant antibiotics are highly repeatable. (**a**) Experimental design for high-throughput antibiotic resistance evolution. An automated feedback loop keeps each individual bacterial population growing in exponential phase at 50% growth inhibition^26^ (Methods). We performed evolution experiments for 258 *E. coli* gene-deletion strains^36^ in triplicate and 36 replicates of a reference strain (*ΔlacA*) under three clinically relevant antibiotics. **(b–d)** Optical density (OD), antibiotic concentration (IC₅₀), and growth rate normalized to that of the reference strain in the absence of drug (*g*/*g*_0_) versus time for twelve replicate populations of the reference strain under NIT selection. Population size (proportional to OD) and selection pressure (quantified by *g*/*g*_0_) are tightly controlled. **(e–g)** IC₅₀ of the reference strain versus time for MEC (e), NIT (f), and TMP (g). The black line shows the median, the light gray area 90%, and the dark gray area 50% of the data, respectively. The blue line shows the variability of the IC₅₀ over time, calculated as the interquartile range normalized by the final fold-increase of the median in logarithmic space (Methods).

To systematically study the effects of the initial genotype on the repeatability of resistance evolution and to mimic spontaneous loss-of-function mutations, we evolved 258 *E. coli* K-12 gene-deletion strains^36^ in three replicates under three different antibiotics for up to three weeks (Fig. 1). To detect any global effects of initial resistance on evolution, if they exist, we selected the 258 gene deletions to cover a wide range of initial resistance levels to the three antibiotics considered in this study, while also representing diverse cellular functions (Fig. S3; Methods). In this assay, we also included 36 replicates of a reference strain (*ΔlacA*). We used *ΔlacA* as a reference because *lacA* is not involved in resistance and *ΔlacA* carries the kanamycin resistance cassette, which is also present in the 258 gene-deletion strains. This makes it more comparable to those than a wild type without any gene deletion (Methods). We selected antibiotics to represent different modes of action: nitrofurantoin (NIT), a prodrug that acts through the formation of nitrogen radicals in the cell^37^; mecillinam (MEC), a beta-lactam antibiotic that targets bacterial cell-wall synthesis^38^; and trimethoprim (TMP), a competitive inhibitor of FolA, a key enzyme in folic-acid synthesis^39^. All three antibiotics are clinically relevant, especially for treating urinary-tract infections (UTIs), which are predominantly caused by *E. coli*^40^. In short, our experiments provide accurate time-resolved measurements of resistance (Fig. 1b–d), allowing quantitative analysis of the phenotypic repeatability of resistance evolution.

### Phenotypic trajectories of resistance evolution to clinically relevant antibiotics are highly repeatable despite inherent stochasticity

We observed rapid resistance evolution to all three antibiotics, although the rate and extent of the resistance increase varied. On average, the resistance of the reference strain increased by at least two orders of magnitude within two to three weeks (Fig. 1e–g). For NIT, most populations approached the solubility limit of the drug within 250 hours, corresponding to a 70-fold increase. TMP resistance increased by 300-fold during the experiment, while MEC resistance showed the highest increase at 390-fold (Fig. 1e–g, Fig. S4a–c). While the rate of resistance increase slowed over the course of the experiment, resistance to TMP and MEC continued to increase until the end of the experiment. Overall, the observed trajectories of resistance gain were characteristic for each antibiotic.

Phenotypic repeatability of resistance evolution was high and strongly dependent on the antibiotic used. The resistance levels of the 36 initially clonal replicates of the reference strain began to diverge for all antibiotics after a few days (Fig. 1e–g), as expected due to the inherent stochasticity in the evolutionary process. This increase in phenotypic divergence was more pronounced for TMP and MEC, but remained small compared to the overall resistance gain (variability in Fig. 1e,g; Fig. S4a,c). In contrast, for NIT, the trajectories of resistance gain were highly parallel and the vast majority remained within a narrow band (Fig. 1f; Fig. S4b). Although the solubility limit compresses late-stage variation for NIT, this result remains valid regardless of this effect (Methods). These observations suggest that while evolutionary stochasticity has a stronger effect for MEC and TMP, the underlying evolutionary dynamics for NIT are nearly deterministic, making phenotypic resistance evolution highly predictable – at least for initially genetically identical bacteria.

### Genetic background can strongly influence phenotypic resistance evolution in ways not explained by global epistasis

To address how the genetic background affects the repeatability of resistance evolution, we analyzed the evolutionary trajectories of the 258 different gene-deletion strains (run in triplicate). Compared to the reference strain, the distributions of resistance gains for the gene-deletion strains at the end of the experiment had a slightly lower median, while, for MEC and TMP, they had a similar width (Fig. 2a). For NIT, the differences were more pronounced: the distribution of resistance gains was broader than for the reference strain, with many gene deletions showing clearly suppressed resistance evolution (Fig. 2a). This divergence emerged abruptly after several days and remained for the remainder of the experiment (Fig. 2c). In contrast, for MEC and TMP, the evolutionary trajectories diverged slowly but continuously throughout the experiment (Fig. 2b,d). For TMP and MEC, evolutionary divergence is only slightly, yet significantly, higher between gene-deletion strains than within replicates of a single strain, i.e., when arising solely from the stochasticity of evolution (Fig. S4a,c). For NIT, this difference is more pronounced (Fig. S4b). These data show that, globally, the effects of the genetic background on phenotypic resistance evolution are moderate for TMP and MEC, but stronger for NIT.

**Fig. 2:**
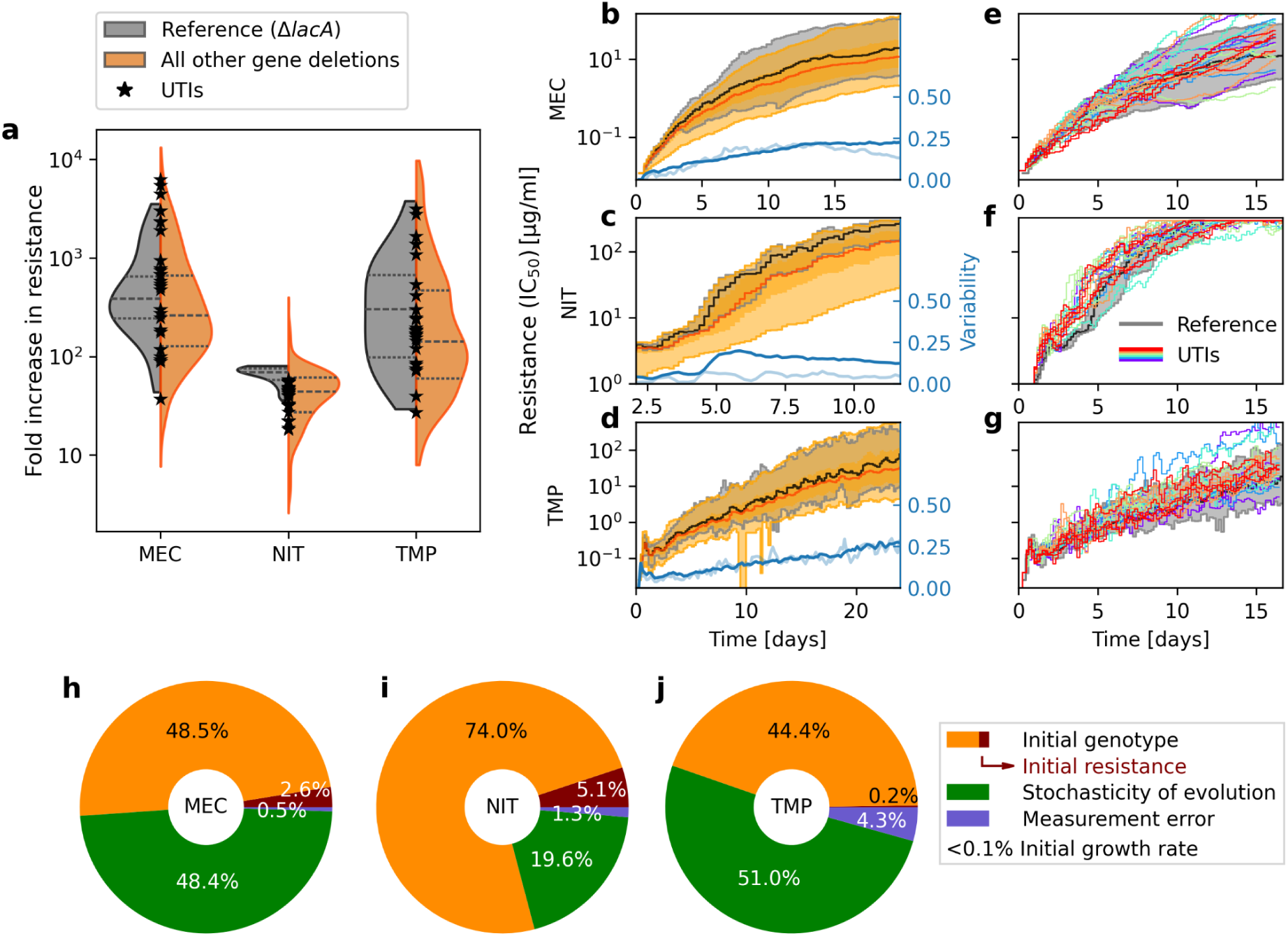
Genetic background can strongly influence phenotypic resistance evolution, in ways not explained by global epistasis. (**a**) Distributions of the final fold increase in resistance (IC₅₀) for the reference strain (gray) and across gene-deletion strains (orange), and values of the same for UTI strains (black stars), for MEC, NIT and TMP (Methods). Inner dashed lines indicate quartiles of the data. **(b–d)** Resistance gain (IC₅₀ versus time) for gene-deletion strains (orange) compared to the reference strain (black). Solid lines show the median, shaded areas show 90% of the data. In blue: variability (as in Fig. 1) within the gene deletions (dark) and within replicate populations for the reference strain (dimmed). **(e–g)** As (b–d), but for different *E. coli* UTI isolates (colored individual trajectories) instead of gene-deletion strains. **(h–j)** Fractions of resistance gain explained by initial resistance, initial growth rate, other characteristics of the initial genotype, evolutionary stochasticity, and measurement error estimated using a linear model that subsequently incorporates these contributions in that order (Methods). Initial resistance generally explains 5% or less of the variance and initial growth rate virtually none (Methods; Fig. S5), showing that these results are not due to common global epistasis.

Resistance evolution remained largely repeatable at the phenotypic level, even for completely different strains of the same species. We performed evolution experiments as before, but using seven different *E. coli* isolates from UTIs^41,42^, each in two or four replicates. Such different *E. coli* isolates typically share less than 75% of their genes^43^. Therefore, their genomes are much more diverse than those of the gene-deletion strains. Despite these substantially different genetic backgrounds, the trajectories of resistance gain were remarkably similar to those observed in the K-12 reference strain (Fig. 2e–g; Fig. S4d–f). Although the evolutionary trajectories of UTI isolates are not as similar to those of the K-12 reference strain as those of other UTI isolates, the differences are much smaller than the total increase in resistance over evolutionary time (Fig. S4d–f). Taken together, these results show that genetic background influences spontaneous antibiotic resistance evolution, but often to a limited extent, allowing lab strains to serve as valuable proxies for clinical isolates in the study of evolutionary dynamics.

The effects of genetic background on resistance evolution were mostly not explained by global epistasis (Fig. 2h–j and Fig. S5). Given the general importance of global epistasis in microbial evolution^8,15,12,9^ and its recent observation in antibiotic resistance evolution experiments^7,26^, we hypothesized that the effects on resistance gains during evolution might be largely determined by the initial resistance of the gene-deletion strains. Initial resistance to all three antibiotics varied by more than an order of magnitude, but it showed little correlation with the resistance gain at the end of the evolution experiment (Fig. S5): Even for NIT, this correlation was weak, despite its artificial amplification due to some populations reaching the solubility limit of the drug (Fig. S5; Pearson’s *r*=-0.23); for MEC, the correlation was also weak and for TMP it was almost nonexistent (Fig. S5; *r*=-0.23 and *r*=-0.08, respectively). Therefore, the variability in resistance gain between different genetic backgrounds cannot be primarily attributed to differences in initial resistance. This is consistent with some recent observations^19,33,44,45^, but in contrast to others^26,46^ (Discussion).

Due to the unexpectedly low explanatory power of global epistasis, we investigated whether factors other than initial resistance could account for the observed resistance gain. We used a linear statistical model to partition the variance into contributions from initial genetic background, evolutionary stochasticity, and measurement error^8^ (Methods). Nearly half of the resistance gain at the end of the experiment was explained by the genetic background, for MEC (51.1%) and TMP (44.6%). Initial resistance accounted for only a small fraction of this contribution. Most of the rest (48.4% and 51.0%, respectively) was attributed to evolutionary stochasticity (Fig. 2h,j). For NIT, the contribution of the genetic background was even higher (79.1%), while the role of evolutionary stochasticity was reduced to 19.6% (Fig. 2i). Taken together with the previous results (Fig. 1f), a consistent picture emerges in which NIT resistance evolution is highly deterministic, but more strongly affected by initial genotype compared to other antibiotics. We conclude that, across antibiotics, genetic background is a crucial factor influencing differences in the ability of bacteria to evolve resistance, while the inherent stochasticity of the evolutionary process imposes clear limits on its predictability.

Similar to initial resistance, initial fitness had a negligible effect on the ability of the gene-deletion strains to evolve resistance. Further partitioning of the contribution of genetic background to the variance in resistance gain into initial resistance, initial fitness (i.e., growth rate in drug-free growth medium), and other genetic background characteristics (Methods) revealed that initial resistance was a minor factor for MEC and TMP, explaining only 2.6% and 0.2% of the variance, respectively, and slightly more (5.1%) for NIT (Fig. 2h–j). Initial fitness had a negligible effect, explaining less than 0.01% of the variance for all antibiotics. The latter reassuringly reflects that only strains with minimal fitness defects in the absence of drugs were included in our selection of gene deletions (Methods).

Taken together, the genetic background can have a major impact on the ability to evolve resistance, in a manner that is mostly consistent with idiosyncratic epistasis – rather than the global epistasis commonly observed in microbial evolution experiments. In other words, as-yet-unknown consequences of the gene deletions, rather than their effects on initial resistance and fitness, are at the core of the resistance gains that the bacteria can achieve.

### Fixed mutations are highly convergent at the gene level

Is the high repeatability of phenotypic resistance evolution accompanied by a similar repeatability of the mutations fixed in the populations during the evolution experiment? To explore the relationship between the observed phenotypic resistance gains and the underlying genetic changes, we performed whole-genome sequencing of 316 selected populations before and after evolution (Methods). For each antibiotic, the selection included 5 to 8 replicates of the reference strain and all three evolutionary replicates for each of at least 30 gene-deletion strains whose resistance trajectories clearly diverged from the reference strain (Table S1; Methods). Note that this selection biases the sample toward lower repeatability. Some populations exhibited a mutator phenotype that spontaneously emerged during the experiment, as evidenced by a much larger number of fixed mutations than in most samples. Known mutators (e.g., *ΔmutL*) displayed this phenotype from the outset of the experiment. To systematically identify all mutator phenotypes, we performed permutation tests in which we recursively removed samples with a greater number of mutations than expected under the null model where all populations were equally likely to acquire mutations (Methods). Twenty mutator populations identified in this way (none for NIT, 6 for MEC and 14 for TMP; Table S2) were excluded from further analysis. The remaining 269 populations had 385 mutations for NIT, 443 for MEC, and 600 for TMP by the end of the experiment. Most of the mutations occurred in coding sequences and were nonsynonymous (81.4%), while a smaller percentage (11.4%) occurred in promoter regions (Fig. S6). We mapped each putatively functional mutation to a gene or promoter region, and then analyzed repeatability at the gene level.

Resistance mutations showed high convergence at the gene level. For all three antibiotics, the most frequently mutated gene was mutated in up to 68% of the populations, and the second most frequently mutated gene in over 50% (Fig. 3a). Overall, NIT had the lowest diversity of mutated genes, despite the relatively strong effects of genetic background on phenotypic resistance trajectories for this drug (Fig. 2i); repeatability was lower for MEC and TMP (Fig. 3a, inset). We quantified the degree of repeatability in mutated genes through the convergence index (CI), which was significantly higher than expected by chance (*p*=10^-4^ for all three antibiotics, Methods). The high mutation repeatability indicates that most gene deletions have a limited effect on the mutational path to antibiotic resistance.

**Fig. 3:**
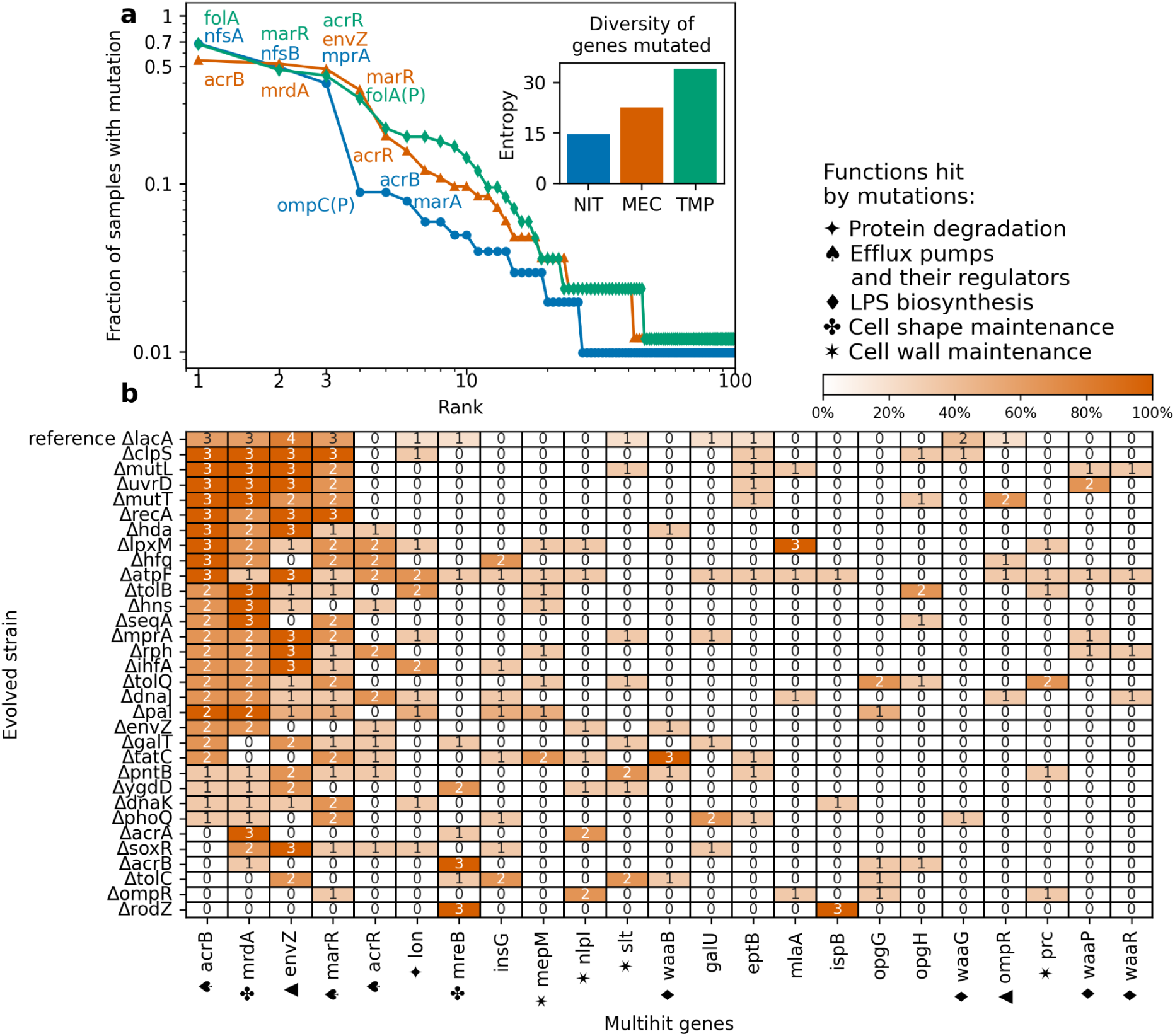
Fixed mutations are highly convergent at the gene level. (**a**) Rank-ordered mutated genes (*x* axis) and the fraction of evolved, sequenced populations showing at least one mutation in those genes (*y* axis). Rank one refers to the most mutated gene across all sequenced populations for each antibiotic, rank two to the second most mutated, and so on. The names of some genes up to rank 6 are shown for each antibiotic. Inset shows the entropy as a measure of mutation diversity for each antibiotic. **(b)** Overview of multihit-gene mutations fixed in all sequenced evolved populations for MEC (Methods; TMP and NIT in Fig. S8). Multihit genes are shown rank-ordered. Numbers and color intensity indicate the number and percentage of replicate populations with the mutation, respectively.

The specific genes that were hit by mutations in more samples than expected by chance (i.e., *multihit genes*, Fig. 3b and Fig. S8; Methods), likely contribute to the evolved resistance to each antibiotic. The most frequently mutated genes were known resistance targets often involved in the first steps toward resistance, in particular *folA*, encoding the target of TMP, and *nfsA*, an enzyme that activates the prodrug NIT. Notably, genes encoding components of multidrug efflux pumps (*acrB*) and their regulators (*marR*, *mprA*) were among the most frequently mutated for all three antibiotics, generalizing previous observations for other antibiotics^31,26,47^. For MEC, the efflux-pump component *acrB* was mutated even more frequently than *mrdA*, which encodes the target of this drug (Fig. 3). In short, for all three antibiotics, evolution consistently converged on mutations affecting both the respective drug targets and efflux pumps.

### Specific cellular functions are general modifiers of resistance evolvability, while others have antibiotic-specific effects

Despite the overall high repeatability of both phenotypic trajectories and mutational paths across different genetic backgrounds, we identified several instances among the 258 gene-deletion strains where evolution produced qualitatively different outcomes. We systematically detected cases where the resistance trajectories clearly deviated from those observed in the reference strain (Fig. 1e–g) and ranked them based on the magnitude of this deviation (Methods). Strains with drastically increased mutation rate (e.g., *ΔmutL*) often accelerated resistance evolution, as expected^48^ (Fig. 4a; Fig. S9–S10). Notably, disrupting specific cellular functions consistently slowed resistance evolution across different antibiotics. Deletions of components and regulators of multidrug efflux pumps (e.g., *acrB* and *tolC*)^49^ drastically reduced resistance evolution for all three antibiotics tested here (Fig. 4; Fig. S9–S11), as well as for tetracycline and chloramphenicol^26^. Similarly, deletions of global regulators (*hns*, *fis*), chaperones (*dnaK*), and genes involved in lipopolysaccharide (LPS) biosynthesis (*lpxM*) generally slowed resistance evolution (Fig. 4; Fig. S9–S11). Our experimental design and controls ensure that the effects of gene deletions are not simply due to changes in initial drug susceptibility or population size (which is tightly controlled, Fig. 1a-d), growth defects (shown to have negligible effects, Fig. 2h–j), or contamination (excluded by media-only controls and whole-genome sequencing; Methods). Together with other recent work^33,50^ (Discussion), these results suggest that a limited number of specific cellular functions act as general modifiers of resistance evolvability, effective across antibiotics with distinct modes of action. In the following, we call this effect *function-specific epistasis* (Fig. S1c).

**Fig. 4:**
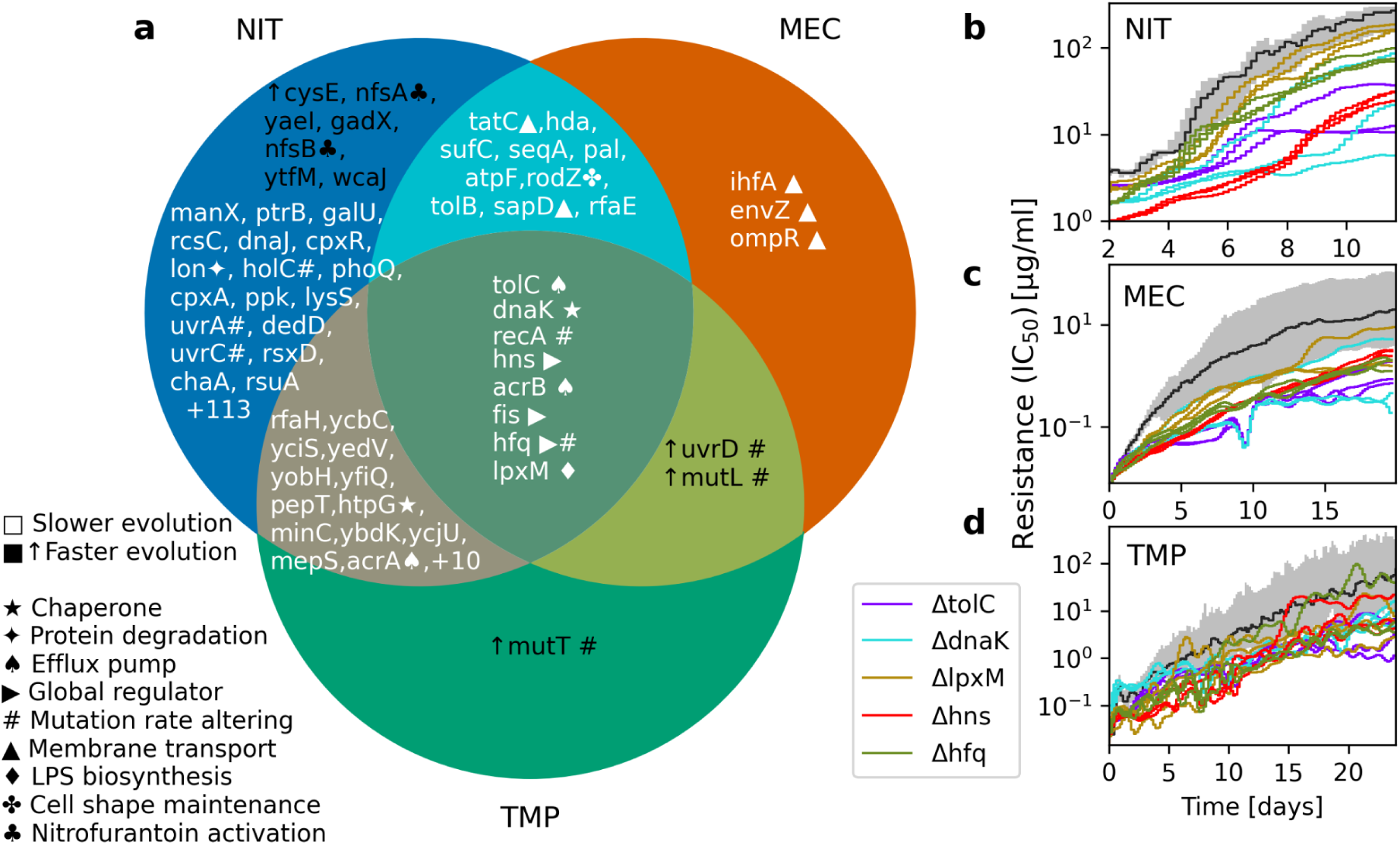
Specific cellular functions are general modifiers of resistance evolvability, while others have antibiotic-specific effects. (**a**) Venn diagram of gene deletions that significantly alter resistance evolvability for NIT (blue), MEC (orange), and TMP (green) compared to the reference strain (Mann–Whitney *U* test on a score based on the area between the IC₅₀ time series, *p*<0.05 after false discovery control; Methods; Fig. S9–Fig. S11). Gene deletions are listed in order of decreasing significance for the respective antibiotic(s). Gene deletions shown in black with an upward arrow (↑) accelerate resistance evolution, while all others slow resistance evolution. **(b–d)** IC₅₀ versus time for three replicates of selected gene-deletion strains, each representing one of the functions in the center in (a) (smoothed individual trajectories, Methods) and for the reference strain (as in Fig. 1e–g).

In addition to cellular functions that modify resistance evolvability in general, we identified others that have specific effects for individual antibiotics. For NIT, these included protein degradation (*lon*, *clpS*), regulation of chromosome-replication initiation (*seqA*, *hda*, *dam*), response to oxidative stress (*azoR*, *ahpC*, *gntY*), and response to DNA damage (*uvrA*, *uvrC*, *holC*) (Fig. 4a; Fig. S11). For MEC, disruption of several genes affecting the function or expression of the outer membrane porins OmpC and OmpF, required for antibiotic uptake (*envZ, ompR, sapD*) slowed down resistance evolution (Fig. 4a; Fig. S9). Disruption of the cytoskeletal protein RodZ, which acts to maintain cell shape, had a similar effect, likely because it also reduces expression of *ompC* and *ompF*^51^. Apart from the general modifiers mentioned above, only one unique gene deletion for TMP (*mutT*, a mutator) showed a significant deviation from the resistance trajectories in the reference strain (Fig. 4a; Fig. S10). This might be partly because TMP resistance often arises from mutations that alter the structure and expression level of the drug target FolA (Fig. 3a) – a path to resistance that may be largely independent of genetic perturbations elsewhere in the genome.

### Function-specific epistasis can substantially alter mutational paths to resistance

While the vast majority of gene-deletion strains carry beneficial mutations in the same commonly hit genes, a few result in fundamentally altered mutational paths (Fig. 3; Fig. S8). To determine whether genetic background generally affects the mutated genes, we analyzed whether mutations in evolutionary replicates of the same deletion strain were more likely to hit the same genes than those in unrelated strains. We used the parallelism index (*PI*) as a quantitative measure of such parallel evolution^8^ and compared the observed hit genes in the gene-deletion strains with a randomly reshuffled ensemble, which destroys any relationship with the specific gene deletion but preserves the remaining structure in the data (Methods). This global analysis revealed that the genetic background had a significant effect on the selected mutations for all three antibiotics (Fig. 5a–c; *p*=0.0068 for NIT, *p*=10^−4^ for MEC, *p*=0.048 for TMP). Thus, single-gene deletions alone are capable of qualitatively altering the mutational paths to antibiotic resistance.

**Fig. 5:**
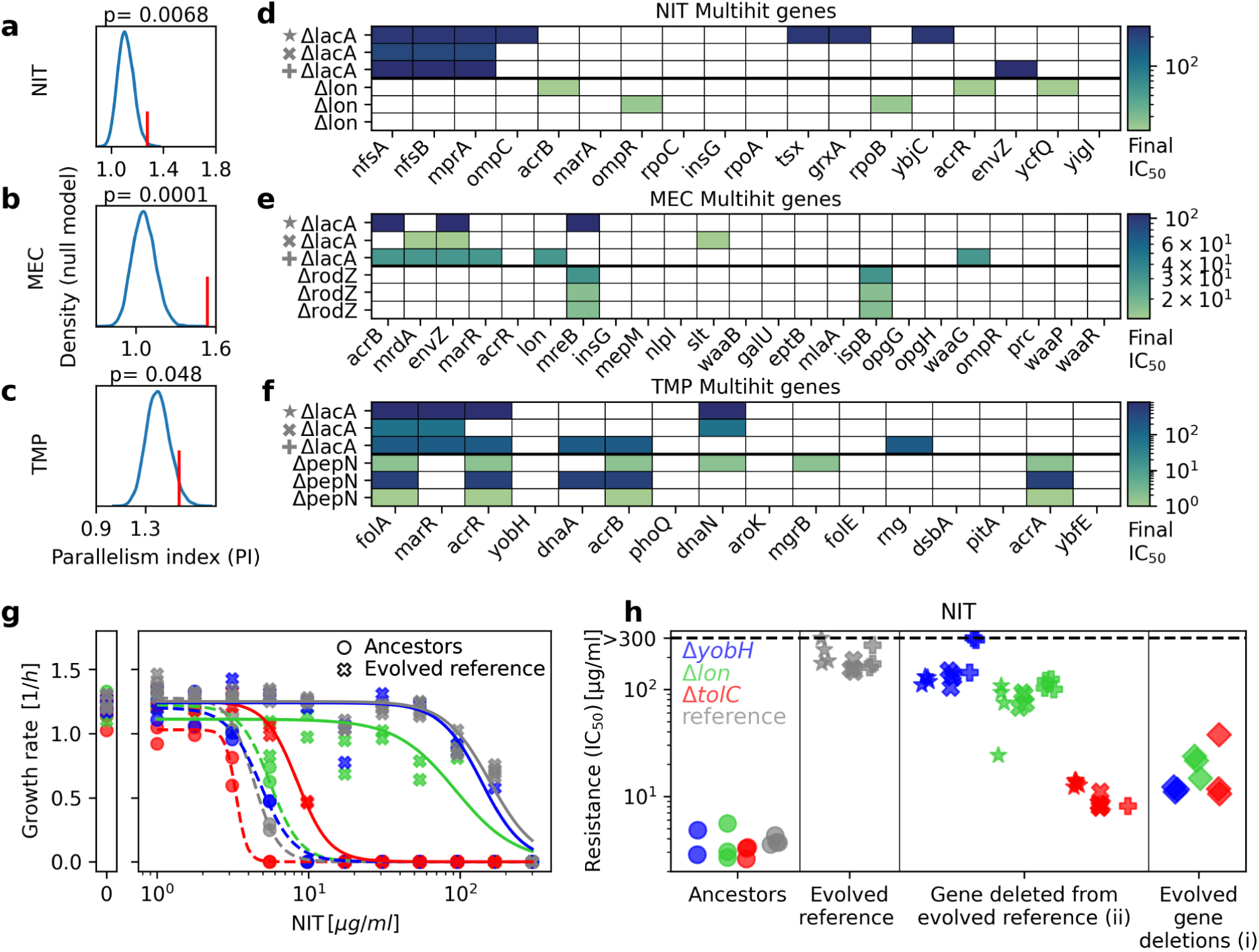
Gene deletions can significantly alter mutational paths to resistance. (**a–c**) Observed parallelism index (red vertical line) compared with a kernel density estimation (blue) of the distribution resulting from a permutation null model in which the gene-deletion information is lost^8^ (Methods). **(d–f)** Fixed mutations in example gene-deletion populations evolved under NIT (d), MEC (e), and TMP (f). Three replicate populations of the *ΔlacA*-evolved reference strain are shown above for comparison (gray symbols indicate the same populations in panel h). Colored squares indicate mutated genes, and the color indicates the final IC₅₀ of the evolved population; blank squares indicate loci that were not mutated. Genes are rank-ordered by number of hits, as in Fig. S8 and Fig. 3a. No additional genes are mutated for *Δlon* and *ΔrodZ*. The complete set of mutations in higher-rank genes for *ΔpepN* is shown in Table S1. **(g–h)** Comparison of (i) the extent to which lineages starting with a gene deletion evolve lower levels of resistance, versus (ii) the extent to which resistance drops when these same genes are deleted from a *ΔlacA*-evolved reference population (shown here for NIT; data for TMP and MEC are in Fig. S12). **(g)** Examples of dose-response curves for ancestors (circles) and evolved reference strains (crosses) with and without one additional gene deletion (*Δlon*, green; *ΔtolC*, red; *ΔyobH*, blue). Repeated symbols show technical replicates used for the estimation of the dose-response parameters. **(h)** Comparison of the IC₅₀ values obtained from (g). Colors represent different gene deletions, and symbols indicate the evolved reference clones in which the gene deletions were introduced (Methods; *ΔlacA* populations in (d) and Table S1 show the mutations in the evolved reference populations). Black dashed lines indicate the maximum concentration used in the dose-response curve, which is the upper bound for the IC₅₀ detection.

To investigate whether the gene deletions that evolved resistance more slowly (Fig. 4) had altered mutational paths to resistance, we compared their mutation profiles to the most commonly mutated genes, as exemplified by those in the reference strain (Fig. 5d–f). Resistance may evolve more slowly in a different genetic background where common resistance mutations still occur but have weaker beneficial effects due to negative epistasis. Alternatively, the genetic background may pose a more fundamental barrier to resistance, i.e., the common resistance mutations may no longer be effective, thus forcing evolution to explore alternative – typically inferior – mutational paths to resistance.

Although some of the gene deletions that slow resistance evolution carry mutations in the most commonly mutated genes, we observed the most dramatic effects of function-specific epistasis in a handful of gene-deletion strains, where none of the most commonly hit genes were mutated. Notably, for NIT, disruption of the Lon protease consistently prevented mutations in the four most frequently mutated genes from fixing, including the otherwise nearly ubiquitous loss-of-function mutations in *nfsA* and *nfsB*, which hinder activation of this prodrug (Fig. 5d; Fig. S8). We were able to generate a viable strain with both *lon* and *nfsA* deleted, indicating that this phenomenon is not due to synthetic lethality. Notably,

loss-of-function mutations in *lon* were previously proposed to limit the evolvability of a cryophile adapting to elevated temperatures^52^. For MEC, the *rodZ* deletion consistently prevented mutations in all six of the most common multihit genes (Fig. 5e; Fig. 3b). No such drastic effects occurred for TMP: mutations affecting the most commonly mutated gene *folA* were not prevented by any gene deletions. However, the second and third most common multihit genes, which affect the expression of the AcrAB–TolC multidrug efflux pump, were no longer fixed in several deletion strains that disrupt the function of these pumps (Δ*tolC*, Δ*acrA* in Fig. S8). These examples and others (Fig. S8) underscore the potential of targeting individual genes to not only slow resistance evolution, but also to fundamentally disrupt the underlying evolutionary dynamics.

To determine the cause of these unexpected effects on evolutionary dynamics, we investigated whether strong genetic interactions could explain the observed phenomena. In this case, the usual resistance mutations would no longer be beneficial in the absence of the relevant genes (Fig. 4a). Thus, we hypothesized that deleting the specific genes from evolved strains carrying those resistance mutations would reduce their resistance. We expected their resistance levels to drop to values similar to those of populations that evolved from ancestors already containing the specific gene deletion. To test this hypothesis, we examined three clones of the *ΔlacA*-evolved reference strain that harbor resistance mutations in common multihit genes (Fig. 3a). These three clones were isolated from the endpoint populations of three independent evolution experiments for each antibiotic (Methods). We deleted genes affecting mutational paths (*tolC, lon, rodZ, waaP, pepN, ompR,* and *yobH*) from these clones (Fig. 5g–h; Fig. S12). We assessed the effect of the gene deletions on antibiotic susceptibility using dose-response curve measurements (Fig. 5g; Fig. S12). Only in the case of the efflux-pump disruption Δ*tolC* did deleting the gene from the *ΔlacA*-evolved background greatly reduce the effect of the common resistance mutations. For NIT, the IC₅₀ decreased to levels similar to those of the evolved gene-deletion strain, as hypothesized (Fig. 5g–h). For MEC and TMP, deleting *tolC* in the evolved reference strain background slightly reduced the IC₅₀ as well (Fig. S12). This strong negative epistasis observed for NIT is consistent with previous observations^26^. These results show that specific gene deletions can exhibit strong negative epistatic interactions with the common resistance mutations, effectively eliminating their beneficial effects and forcing evolution to explore alternative paths to resistance.

Unexpectedly, however, the majority of gene deletions (6 out of 7) had little effect on the evolved resistance: here, the common resistance mutations conferred almost the same increase in IC₅₀ in the deletion background as in the reference strain (Fig. 5h and S12). Notable examples of this phenomenon are the deletions of *lon*, *rodZ*, and *pepN*, which left evolved NIT, MEC and TMP resistance, respectively, almost unchanged (Fig. 5h and S12). This suggests that strong negative epistasis is the exception rather than the rule: often, the observed function-specific epistasis cannot be directly explained by negative epistatic interactions. Further elucidating the underlying molecular mechanisms can be difficult, especially for genes with promiscuous intracellular interactions and pleiotropic effects, such as *lon*, and is beyond the scope of this study (Discussion).

### Small-molecule inhibitors of the identified cellular targets may affect resistance evolution

Function-specific epistasis could potentially be leveraged to impede resistance evolution. Such an intervention strategy would combine antibiotics with small molecule inhibitors of the identified targets (Fig. 4) if such inhibitors exist or can be developed. Motivated by the broad effects of efflux pumps and chaperones on resistance evolution, we attempted to partially mimic the effects of the corresponding gene deletions (*tolC*, *dnaK*) with small-molecule inhibitors. Drugs against these targets already exist: DnaK was recently shown to be inhibited by telaprevir, an FDA-approved hepatitis C drug^35^; the AcrAB–TolC pump is inhibited by domperidone^53^, a dopamine antagonist used to treat nausea. We performed an evolution experiment as before (Fig. 1), but in the presence of these small molecules at high concentrations that did not affect growth (Methods). In addition to the reference strain, we included seven *E. coli* isolates from UTIs^41,42^ and deletion strains directly related to multidrug efflux pumps (e.g., *acrB*) in this experiment.

The small-molecule inhibitors affected resistance evolution (Fig. 6; Fig. S13, Methods). Notably, the efflux-pump inhibitor domperidone had a significant global tendency to slow down MEC resistance evolution for UTI isolates and the reference strain (Fig. 6d; combined *p* = 0.01; Methods), as predicted from the effects of deleting efflux-pump components (Fig. 4). This tendency was not observed for TMP (Fig. S13), where the deletion of efflux-pump components had a significant, albeit smaller, effect (Fig. 4d; Fig. S10). This smaller effect size is likely due to the fact that resistance to this drug primarily evolves through mutations affecting the drug target FolA. At the level of individual strains, domperidone strongly affected the evolution of Δ*acrA* and Δ*acrB* to MEC and NIT (Fig. 6d; Fig. S13). This suggests that domperidone and the *acrA* or *acrB* deletion both partially impair efflux-pump function, but together they synergistically disable the efflux pump, thereby blocking this common path to resistance, similar to the effect of the *tolC* deletion. Note that this interpretation would imply that domperidone does not inhibit the efflux pump by binding to AcrB alone, contrary to computational predictions^53^. The DnaK*-*inhibitor telaprevir had no significant global effect on resistance evolution (Fig. S13; Methods). Unexpectedly, it had opposite effects on different strains: it slowed resistance evolution for some and accelerated it for others (Fig. S13). This suggests a complex role for chaperones in evolution, consistent with previous reports indicating opposite effects of chaperone perturbations on evolution in other contexts^54–57^. Overall, these data indicate that, with further optimization, drugs could be developed to slow antibiotic resistance in ways that are partially predictable based on the effects of genetic perturbations on evolution.

**Fig. 6:**
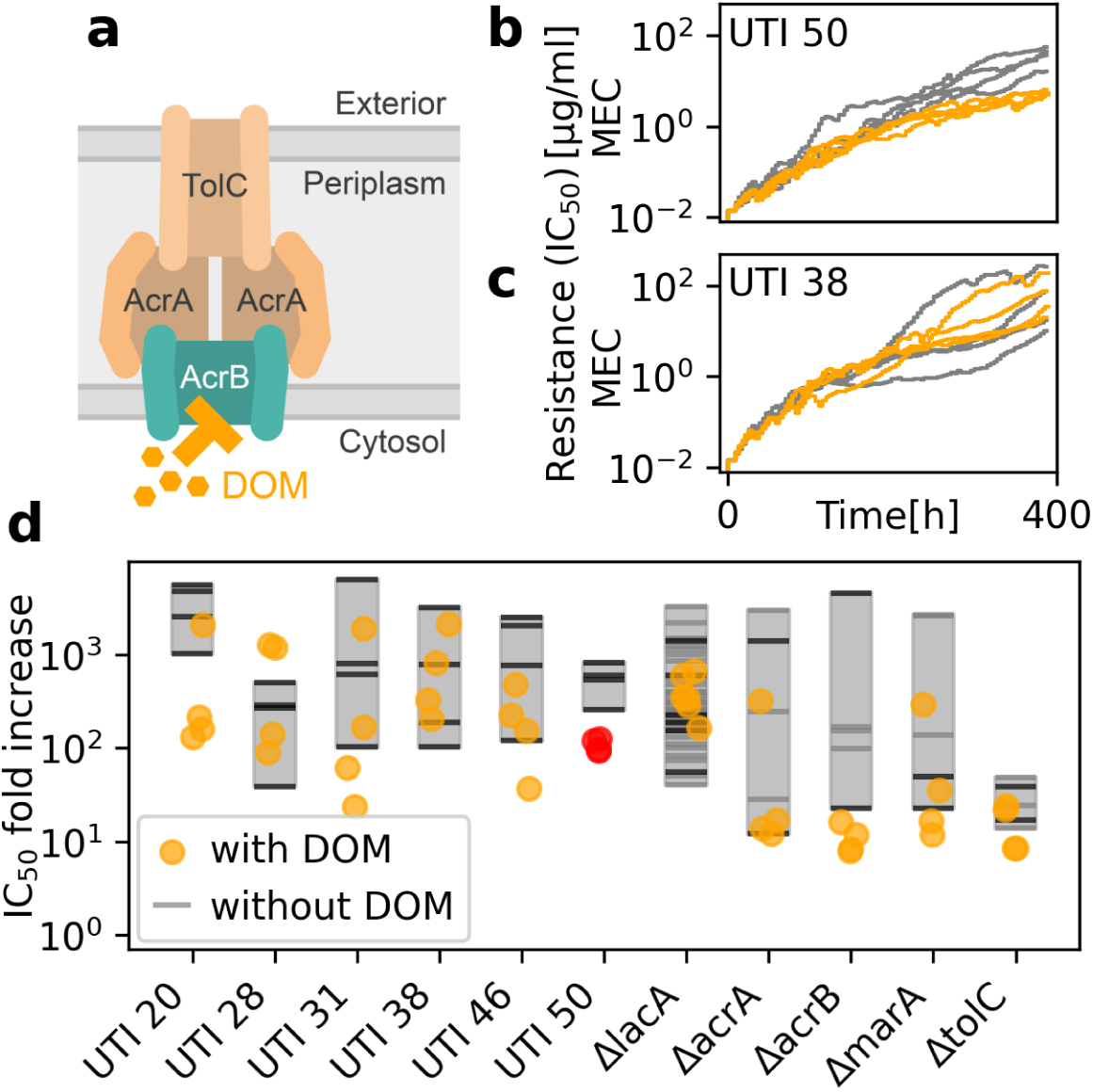
Function-specific epistasis can be exploited to slow resistance evolution. (**a**) Schematic of the AcrAB–TolC efflux pump, highlighting the small-molecule inhibitor domperidone (DOM). **(b–c)** Resistance-increase trajectories of two exemplary evolving UTI isolates in the presence (orange lines) or absence (gray lines) of domperidone. **(d)** Comparison of IC₅₀ fold increase in the presence or absence of domperidone for different strains (shown here for MEC, full data set in Fig. S13). Gray and black lines show replicates of the reference strain without domperidone from two different evolution experiments (MEC_evo and inhibitors_evo, respectively, in Table 1). Gray rectangles show the range of fold increase for easier comparison; orange dots show replicates evolved with domperidone without a significant change; red dots indicate that domperidone had a significant effect on UTI 50 (*p*=0.043, two-sample *t*-test, false-discovery rate controlled with Benjamini–Hochberg adjustment). The global trend of reduced resistance evolution observed with domperidone is significant (combined *p* = 0.01, Methods).

**Table 1:** Summary of evolution experiments and corresponding parameters. **Steepness** (*n*) was estimated as the exponent of a Hill function fit (Methods), averaged over several typical dose-response curves of ancestors. The “maximum update factor” is an upper bound on the calculated factor by which the antibiotic concentration increases with each dilution, which is different for the different antibiotics, depending on the steepness of their dose-response curves (Methods).

| Experiment name | Antibiotic | Duration [days] | [total # of dilutions] | # of plates | Contami -nations | Steepness ($n$) | Maximum update factor | Initial concentration [$\mu\text{g/ml}$] | Maximum concentration [$\mu\text{g/ml}$] |
| --- | --- | --- | --- | --- | --- | --- | --- | --- | --- |
| **NIT_evo** | NIT | 12 | 63 | 9 | 1/54 | 4.39 | 5 | 0.3 | 300 (solubility limit) |
| **TMP_evo** | TMP | 24 | 129 | 9 | 9/54 | 1.3 | 5 | 0.025 | 1500 |
| **MEC_evo** | MEC | 20 | 107 | 9 | 0/54 | 7.74 | 1.34 | 0.0115 | 300 |
| **inhibitors_evo** | NIT, TMP, MEC + domperidone or telaprevir | 16 | 96 | 6 | 2/24 | MEC 7.74<br>NIT 3.8<br>TMP 1.53 | $9.5^{1/n}$ | MEC 0.014<br>NIT 1<br>TMP 0.04 | NIT: 300,<br>TMP: 950,<br>MEC: 470 |

## Discussion

By quantitatively following the evolutionary trajectories of hundreds of genetically distinct *E. coli* strains to high resistance levels, we found that overall resistance evolution is highly repeatable at both the phenotypic (Fig. 1) and genotypic (Fig. 3) levels, albeit with variation across antibiotics. Our data show that the genetic background strongly influences resistance evolution, accounting for up to three quarters of the variability in resistance gain, with only a small fraction explained by the initial resistance level via global epistasis (Fig. 2). The remaining fraction explained by the genetic background is mostly due to function-specific epistasis: we have identified specific cellular functions, including efflux pumps, chaperones, global regulators, and LPS biosynthesis, that influence resistance evolution across antibiotics (Fig. 4). Resistance evolution to some of the antibiotics in our dataset is also affected by gene deletions involved in other functions, including protein degradation and cell shape maintenance. Disruption of these functions can dramatically alter the otherwise highly repeatable mutational paths toward resistance (Fig. 5). Small-molecule inhibitors targeting the identified functions can induce modest deviations from the otherwise repeatable resistance-increase trajectories (Fig. 6).

A notable difference between our results and several previous studies on microbial evolution is that global epistasis explains only a small fraction of the effect of genotype on resistance gain (Fig. 2h–j; Fig. S5). The prevalence of global epistasis in antibiotic resistance evolution remains debated. Some studies reported patterns consistent with diminishing returns^26,46^, while others found little to no evidence of such global trends^19,33,45^. The reasons for these discrepancies are unclear. One possibility is that the experimental limitations of the different studies prevented the observation of global patterns. For example, in fluctuation assays, where the resolution is limited by the discrete steps in MIC^19,46^, this limitation has been proposed to hinder the detection of global patterns^19^. However, global patterns can be detectable despite this limitation^46^. A more severe limitation may be considering only a relatively small number of genetic backgrounds^19^. In other studies, the resistance increase over the course of the experiments was limited, despite their relatively long duration^33^, which generally makes it harder to detect a global pattern.

Pervasive global epistasis has been observed for fitness^8,15,16^ and similarly for resistance to the antibiotic tetracycline^26^. In evolution experiments, fitness is often quantified as growth rate, and thus can quickly approach an upper bound imposed by fundamental limits on how fast a cell can grow. This effect can be described by a concave nonlinear transformation of a latent additive variable, resulting in a diminishing-returns trend^12,58^. Similarly, resistance to tetracyclines saturates relatively quickly^31^ and tends to converge to a fixed upper bound across genetic backgrounds, resulting in a diminishing-returns trend^26^. However, our data show that such convergence to a fixed resistance level, independent of genetic background, may not occur for most antibiotics. In most cases, the resistance trajectories for the antibiotics studied here continuously increase and diverge, even as they slow down toward the end of the experiment (Fig. 1e–g; Fig. 2b–g). Only for NIT did some convergence of resistance trajectories occur later in the experiment (Fig. 1f), as the solubility limit of this drug was approached. Importantly, our experimental design allowed resistance to increase by several orders of magnitude, bringing populations closer to the regime in which saturation effects might be expected. Despite overcoming several limitations of previous studies, our results show little evidence of global epistasis. This suggests that these patterns are insufficient to explain the effects of genetic background on resistance evolution to the antibiotics considered here.

We found that the effect of different genotypes on the repeatability of evolution is greater for NIT than for MEC and TMP (Fig. 2a–d,h–j): many gene deletions slowed down evolution for NIT (Fig. 2a-b; Fig. 4a). A plausible explanation is that fewer resistance mutations are available for NIT. Our sequencing results showed that resistance mutations hit the same genes in most populations under NIT, whereas the diversity of genes hit by mutations was greater for the other two antibiotics (Fig. 3a), consistent with the limited availability of alternative resistance mutations for NIT. Therefore, when a gene deletion prevents the common resistance mutations, there are no easy alternative paths to resistance for NIT, leading to slower evolution. In contrast, for MEC and TMP, evolution can more easily switch to one of the many alternative mutational paths. A scarcity of available resistance mutations for NIT is also consistent with conclusions from measurements of the distribution of fitness effects (DFE) for these antibiotics^20^. For TMP, the most common resistance mutations affect the expression and function of the drug target FolA^31,59,60^. This likely contributes to making the evolution of TMP resistance even less dependent on genetic background, as most gene deletions do not interact with the drug-target mutations, allowing evolution to proceed unimpeded.

Our finding that disrupting specific functions can slow resistance evolution (Fig. 4) raises the question of the underlying molecular mechanism. The general effect of disrupting multidrug efflux pumps across antibiotics is plausible since these pumps can expel all antibiotics tested^47,49^ and mutations affecting efflux pump function and expression are among the most common for all antibiotics (Fig. 3a). Deletion of an essential component such as *tolC* will render these pumps essentially nonfunctional, thereby forcing evolution to use alternative mutational paths. For MEC, deletion of *tolC* further results in downregulation of the outer membrane porins *ompC* and *ompF*^61,62^, which are required for the uptake of this drug. Broadly consistent with our observations, a general role for chaperones in the evolutionary process has also been proposed, although it is controversial whether chaperone deletions should accelerate or slow evolution^54,55,63^. The role of global regulators and LPS is similarly complex: disruption of either leads to massive perturbations that irreversibly affect global gene expression and cell envelope permeability to small molecules, respectively. This makes it plausible that they can have the strong effects on resistance evolvability we observed. Another study focused on transcription factors and identified additional evolvability suppressors that are common to several other antibiotics^33^. While our study would have excluded some of the relevant gene deletions due to severe growth defects, this suggests that there might be cellular functions, such as drug efflux, protein folding and global regulation, that more generally affect resistance evolution across groups of antibiotics, for which these functions play a direct or indirect role in the resistance mechanisms.

One of the most striking observations we have made is that a few gene deletions (including *lon* and *rodZ*) dramatically alter the mutational path for specific antibiotics, such that none of the most common resistance mutations fix (Fig. 5d,e). Sometimes, these effects can be explained by direct epistatic interactions between the gene deletion and the common resistance mutations, but often they cannot (Fig. 5g–h; S12). In the latter cases, there may be more complex epistatic interactions with intermediate mutational points in the evolutionary pathway towards resistance. Elucidating the underlying molecular mechanisms is challenging due to the promiscuous roles of these genes in the cell. For instance, the Lon protease can degrade various proteins^49^, including regulators that affect the expression of numerous genes, each of which may be crucial for the observed effect. One example is SulA, a component of the DNA damage (SOS) response induced by NIT, which inhibits cell division^64^. SulA protein levels are greatly increased in a *Δlon* background, enhancing NIT toxicity^30^. In addition, Lon degrades the transcription factors MarA and SoxS, both of which activate *nfsA* and *nfsB*^49^. Their expression levels are therefore likely to be increased in *Δlon*, thus facilitating NIT activation and further enhancing its toxicity. These effects may contribute to the slower resistance evolution, but it remains a mystery why the common resistance mutations in *nfsA* and *nfsB* are not selected in this background.

The Lon example illustrates that perturbing broadly acting genes can slow down resistance evolution, probably by reducing access to certain mutational paths or by making alternative routes more costly. Although ancestral growth defects were minimized in our gene selection, disrupting genes with broad roles can still impose general physiological constraints, such as reducing metabolic flexibility, or stress tolerance or perturbing regulatory networks stability. One way to learn more about the underlying mechanisms would be to test mutations specific to the identified gene deletion backgrounds in both those backgrounds and the reference. This could reveal relevant epistatic interactions, offering an exciting direction for future work.

Although we focused on fundamental aspects of evolutionary dynamics, our results have potential long-term implications for antibiotic treatment strategies. It is encouraging that the dynamics of resistance evolution were similar in *E. coli* isolates from UTIs and in the K-12 lab strain (Fig. 2e–g; Fig. S4d–f). In addition, the most commonly mutated genes in our study (Fig. 3a; Fig. S8) are frequently mutated in clinical *E. coli* isolates^65,66,60,59^. The specific genes deleted in the strains we identified (Fig. 4) could serve as targets for new drugs that, when combined with antibiotics, could slow resistance evolution and help maintain the efficacy of existing antibiotics. Our approach revealed a plethora of such targets that affect resistance evolution, complementing previous efforts focused on reducing mutation rates to slow down pathogen evolution by preventing (stress-induced) mutagenesis^67–74^. Particularly promising is the discovery that several targets have a broad impact on the evolution of resistance to multiple antibiotics (Fig. 4). Our finding that the efflux-pump inhibitor domperidone can moderately inhibit resistance evolution in *E. coli* isolates from UTIs (Fig. 6) provides a first glimpse of the potential of this approach. There are a number of other small molecules that have the potential to inhibit efflux pumps^75^. However, further refinement of such small molecules is certainly needed. The development of small-molecule inhibitors of targets identified by our approach and the study of their impact on the evolution of antibiotic resistance is an enticing prospect for future studies.

## Methods

### Experimental procedures

#### Strains, media and drugs

All *E. coli* gene-deletion strains are from the Keio collection and contain a kanamycin cassette instead of the respective gene^36^. To ensure that any differences between the strains are not due to the presence of the cassette, we used the Δ*lacA* strain as a reference because *lacA* is not involved in resistance and the growth medium did not contain lactose. Additional *E. coli* strains included in the experiments with small-molecule inhibitors are isolates from urinary-tract infections in elderly patients selected based on low levels of resistance to the antibiotics used in this study as well as similar growth to the Δ*lacA* reference strain under the following experimental conditions^41,42^. All cultures grew in lysogeny broth (LB) at 30°C (LB agar: Sigma-Aldrich L2897, LB Sigma-Aldrich L3022). This temperature was chosen for technical reasons in the evolution experiment. First, a lower growth rate compared to 37°C is required to maintain exponential growth with the implemented dilution frequency. Second, temperature gradients between the incubator and the climate-controlled room are avoided (section *Automated high-throughput morbidostat evolution experiments*). Since 30°C is not a stressful condition for *E. coli*, and its cell physiology is very similar to that at 37°C^76^, we expect the antibiotic selection pressure to dominate the evolutionary dynamics. While this reduces comparability with studies conducted at 37°C^45,46^ or 34°C^33,48^, it facilitates comparison to our previous results for ribosome-inhibitor antibiotics^50^. For long-term storage, bacterial cultures were supplemented with 15% glycerol and kept at −80°C. Unless otherwise specified, all reagents were obtained from Sigma Aldrich. Catalog numbers: NIT (N7878), TMP (92131), MEC (33447), kanamycin (K4000), ampicillin (2003492, Labomedic), domperidone (D122), telaprevir (ACRO466892500, VWR). Antibiotic stocks were prepared in DMSO (NIT at 30 mg/ml and TMP at 50 mg/ml) or water (MEC at 20 mg/ml, kanamycin at 25 mg/ml, ampicillin at 100 mg/ml). The stocks were sterile-filtered through a 0.22 μm filter and stored in the dark at −20°C. To avoid concentrations of DMSO higher than 3% in the wells, TMP was directly dissolved in LB and sterile-filtered for the highest concentration in the evolution experiment. Small-molecule inhibitor stocks were prepared in DMSO, sterile-filtered and stored in the dark at −20°C (domperidone at 10 mg/ml and telaprevir at 50 mg/ml).

#### Selection of gene-deletion strains for evolution experiments

The 258 gene-deletion strains used as starting points for the evolution experiments were selected from the Keio collection^36^. To address the effects of initial resistance on the final resistance increase, we aimed to select ancestral gene-deletion strains that cover a wide range of initial resistances to the three antibiotics considered in this study. First, we chose 148 gene deletions including all used in^26^ as well as additional genes known to be implicated in antibiotic resistance^20,77,24,31,49^. Then, we randomly selected an additional 110 gene deletions from the subset of 2,225 out of 3,985 Keio collection strains that complied with the following criteria: (a) the initial resistance level was determined in^20^; (b) no growth defect was detected (i.e., *g*_0, *gene del*_. ≥ 0. 8 *g*_0, *reference*_); and (c) the UniProt annotation was sufficiently good (i.e., annotation score greater than three out of five). Note that no strains from KEIO collection Plate 89 were selected as this plate was not in our cryo stocks. A score defined as the mean of the standard deviations of the initial resistance distributions for the three antibiotics was calculated for one full set of 258 pre-chosen plus randomly selected gene deletions. Further optimization of the set to maximize variability was achieved by recursively replacing randomly selected strains and keeping only those that increased the score. Several runs yielded similar score values, and the final set was chosen to have a high score and cover most gene-ontology functions (Fig. S3).

#### Automated high-throughput morbidostat evolution experiments

We implemented a modified version of the high-throughput morbidostat technique from^26,31^ to increase throughput further and enable the use of more diverse antibiotics. In brief, cultures are regularly diluted to ensure exponential growth throughout the entire experiment (Fig. 1). Frequent OD₆₀₀ determination allows observation of growth rates during the experiment. The dilution factor is individually calculated for each culture in each dilution based on an OD measurement right before dilution, ensuring that all cultures are diluted to the same fixed OD value, thereby tightly controlling population size. The antibiotic concentration is updated at each dilution to consistently maintain 50% growth inhibition and sustain selection pressure.

The lab evolution data presented in this manuscript come from four experiments that run for 12 to 24 days (NIT_evo, TMP_evo, MEC_evo, inhibitors_evo in Table 1). Three experiments with single antibiotics (NIT, TMP and MEC) contained the full set of 258 selected gene-deletion strains, for a total of 810 independently evolving populations (experiments NIT_evo, TMP_evo, MEC_evo). The inhibitors experiment (inhibitors_evo) tested multiple conditions, including the three antibiotics and two small-molecule inhibitors, in the same run for a reduced number of gene-deletion strains and additional pathogenic isolates, with a total of 552 parallel populations.

First, three mother plates containing the selected gene-deletion strains were prepared. The strains were restreaked from cryo stocks, and single colonies were used to inoculate the corresponding well of a 96-well plate. For the experiments NIT_evo, TMP_evo and MEC_evo, each mother plate contained one copy of 86 selected gene deletions, four copies of the reference strain, and six media-only wells for contamination control (Fig. 1a). Three replicates of each gene deletion were inoculated from these mother plates using a pin tool (VP 408, V&P Scientific). For the experiment with the small-molecule inhibitors (inhibitors_evo), two mother plates were created (one each for domperidone and telaprevir) with 4 or 5 relevant gene-deletion strains including the reference strain, 7 or 6 UTI strains, and 4 media-only wells. Each plate contains two replicates per strain, for each antibiotic and inhibitor condition. Two replicate plates for each inhibitor and one plate each without inhibitor were inoculated using the pin tool. In all experiments, the cultures initially grew without antibiotics until the first dilution, at which point they were diluted to a preset antibiotic concentration identical for all strains. The morbidostat-like update began at the second dilution onwards, except for MEC, which required an additional dilution to the initial fixed concentration to account for the delayed effect of the antibiotic on growth.

The automated platform used in these experiments consists of a shaking incubator (StoreX STX44, Liconic), a 96-channel pipetting unit capable of pipetting independent volumes in each channel (Lynx LM900, Dynamics Devices), a plate shaker (BioShake 3000elm, QInstruments), a plate reader (Synergy H1, BioTek Instruments), storage for fresh and waste plates (NanoServe, Acell, HighRes Biosolutions), positions to de-lid plates (LidValet, HighRes Biosolutions) and several positions for single plates, all integrated together through a robotic arm (Acell, HighRes Biosolutions) and controlled by Cellario scheduling software V3.4.0.2636 (HighRes Biosolutions) supplemented with custom morbidostat scripts. The entire setup is located inside a hood with HEPA-filtered airflow in a climate-controlled room at 30°C. Cultures were grown in clear, flat-bottom 96-well plates with a total volume of 200 μl. In the experiments NIT_evo,TMP_evo and MEC_evo (Table 1), nine plates were run in parallel, and six plates were run in the inhibitor experiment (inhibitor_evo). The plates were incubated in the incubator (30°C, humidity greater than 95% and shaking speed 500 rpm), taken out for an OD measurement every 24 min with prior shaking on the plate shaker and then diluted in the pipetting unit every 4.2 h on average. After every six transfers, the old plate was incubated until saturation, and a cryo stock of the plate was stored.

During each dilution, the culture was transferred to a new plate and adjusted to a target OD₆₀₀ of 0.01. The antibiotic concentration was updated to maintain growth inhibition of each population at 50% relative to the growth rate of the reference ancestor, *ΔlacA*, in drug-free medium (*g*_0_)^26^. The updated antibiotic concentration was estimated based on the population’s growth rate right before the dilution, using the IC_50_ from a dose-response Hill function. The values of steepness and *g*_0_ in the Hill functions were based on those measured for the *ΔlacA* ancestor prior to the evolution experiments (Methods section *Dose-response curves*)^26^. The current growth rate was estimated in each growth interval before dilution by determining the area under the OD_600_ values in the range of 0.01 and 0.15 and calculating the slope of the triangle with the corresponding area. This estimation is more robust than a linear regression. The maximum increase and decrease in concentration were limited to avoid killing the population, especially when growth rates were above *g*_0_. These factors are linked to the steepness of the Hill function, as it determines the concentration range between no and full inhibition (Table 1). Accordingly, the increases of TMP concentration over evolution exhibit a higher variability due to its shallower dose-response curve, which allows for larger changes in antibiotic concentration to achieve the desired effect on growth.

At each dilution, the calculated volumes of culture, medium without antibiotic, and medium with one of two different antibiotic concentrations were mixed to obtain the desired updated concentration. The two available antibiotic concentrations were chosen to best cover the expected range of updated concentrations and were increased throughout the evolution experiment. In some cases, slowly evolving strains could not keep up with the general increasing trend. In these cases, they were diluted first without any antibiotic and then set to the minimum possible concentration from the antibiotic reservoir. The latter procedure is necessary due to a technical limitation: the need to produce a wide range of antibiotic concentrations to capture different resistance levels with a limited number of reservoirs. It causes the sudden drops (observed when diluting without antibiotic) and the spikes (observed when setting the concentration to the minimum possible) in some of the low-resistance trajectories (Fig. S9–S10). This technical limitation of our experiments affects some antibiotics more than others. It leads to more pronounced variations for antibiotics with large differences in IC₅₀ between fast– and slow-evolving strains because a wider dynamic range of drug concentrations must be covered. For NIT, solubility limits the maximum drug concentration. Consequently, the dynamic range that our reservoirs need to cover is relatively small, and the NIT evolution experiments are less affected by this limitation (Fig. S11). In the inhibitor experiment, one set of medium reservoirs contained 25 µg/ml of domperidone or 70 µg/ml telaprevir. These concentrations were chosen as the highest concentrations that did not cause growth inhibition in the ancestors.

Several preventive measures were taken to avoid contamination, the most crucial of which were utilizing dedicated tip boxes for each plate and several others for media and antibiotics, avoiding mixing by pipetting and placing the lidded plates on the shaker instead, and daily replacement of the reservoirs containing media and antibiotics. To monitor contamination, media-only wells were transferred in each dilution step while adding 30% fresh medium. With these preventive measures, we observed contamination in fewer than 17% of the medium-only wells (maximum 9 of 54 media-only wells in the longest evolution experiment with TMP). Due to the higher medium transfer from the previous plate and the absence of a competing inoculum in the media-only wells, we consider this estimate to represent an upper bound of contamination. This is confirmed by WGS of 316 populations, which revealed only 3 gene deletions that did not match the expected one (0.95%, Table S3). Nine populations (out of a total of 2982) died during the evolution experiments and were excluded from all analyses (Table S4).

#### Dose-response curves

Dose-response curves were obtained from the OD_600_ growth curves across a concentration gradient. Precultures of the respective strains were grown overnight either without antibiotics or, in the case of resistant populations from the evolution experiment, with a low level of the respective antibiotic to reduce the likelihood of resistance reversal. The precultures were diluted 1:1,000 in a concentration gradient in 96-well plates, and their OD_600_ was measured in a plate reader every 20–30 min, depending on the number of plates. The cultures were grown in the concentration gradient for at least 30 h, either directly in the plate reader or in the Liconic incubator integrated into the automated setup of the evolution experiment to measure multiple plates in parallel. Exponential growth rates were used as the response variable, except for MEC dose-response curves, where the nonlinear behavior at intermediate antibiotic concentrations renders the fitted growth rate less informative. In that case, the area under the curve (AUC) was used instead. We used growth rate as the response variable whenever possible because it is a more robust and reproducible measure whose absolute value is easily interpreted. However, the results obtained using either of the two response variables are comparable: the IC₅₀ values estimated using the AUC for NIT and TMP correlate almost perfectly (Pearson correlation coefficient: NIT, 0.98; TMP, 0.99; p-value<10^-50^ for both antibiotics) with those estimated using the growth rate.

Exponential growth rates were fitted using the *LinearRegression* function from Scikit-learn v1.5.2. The OD_600_ data were log-transformed and background-subtracted and the first 20 h of measurements were used. The OD_600_ thresholds for the exponential phase were typically 0.008–0.08 and only consecutively increasing data within this range was fit, requiring a minimum of three data points and at least one above an intermediate threshold (typically 0.025). For MEC, inhibition is not clearly characterized by an exponential growth rate. At inhibitory concentrations, an initial growth phase is followed by decay and potential recovery of the OD_600_ later on. The AUC was calculated within the same time frame, considering the area above the lower OD_600_ threshold using a trapezoid approximation, and normalized by the total time range.

Dose-response curves were fitted using a Hill function, 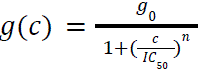, where *g* is the growth rate, *g_0_* is the growth rate without antibiotic, *c* is the antibiotic concentration, *IC_50_* is the concentration at half-maximal inhibition, and *n* is the Hill coefficient. The fit was performed using LMFit v1.3.2 (https://doi.org/10.5281/zenodo.12785036), limiting *g_0_* to two standard deviations around the mean of the measured antibiotic-free growth rates.

#### Gene-deletion mutants of evolved strains

For each antibiotic used in the evolution experiments, we selected three gene deletions to introduce into three different evolved clones of the reference strain (*ΔlacA*). We selected these gene deletions from those that exhibited slower resistance evolution and/or altered mutational pathways (Fig. 4; Fig. 5). To introduce the gene deletions into the evolved *ΔlacA* background, three evolved *ΔlacA* populations per antibiotic were restreaked on LB agar with 50 µg/ml kanamycin. We selected these three populations to best cover the variability in resistance within the 36 evolved *ΔlacA* populations (i.e., one with the highest, one with middle, and one with the lowest resistance). The mutations in all populations can be found in Table S1, where the three selected populations are marked by their symbols in Fig. 5d–f,h and Fig. S12 in a dedicated column named “symbol in Fig. 5 and S12”. Multiple colonies were selected and measured in dose-response curves as described in the previous section. The culture from the closest match to the evolved whole population was chosen.

The FRT-flanked kanamycin cassette in the place of *lacA* was removed via Flp recombination. The evolved *ΔlacA* strains were made competent using a transformation and storage solution (TSS) protocol based on^78^ and the temperature-sensitive plasmid pCP20, which expresses the Flp enzyme, was introduced into each clone. Transformants were selected on LB agar containing 100 µg/ml ampicillin and tested for the absence of growth on 50 µg/ml kanamycin. PCR was used to verify the removal of the FRT-flanked kanamycin cassette. Gene deletions were introduced using standard P1 transduction. KEIO collection strains of the respective gene deletion (i.e., *ΔtolC, ΔrodZ, Δlon, ΔyobH, ΔompR, ΔpepN*, and *ΔwaaP*) were used as donor strains. Overnight cultures of the chosen, evolved *ΔlacA* clones were used as recipients. Colonies were selected on LB agar with kanamycin, and PCR was used to verify the deletion of the target gene in four to eight colonies or all present colonies if there were fewer. Note that in several cases only one successful transformant was isolated. Additionally, the deletion of *rodZ* and *tolC* in MEC-resistant strains was unsuccessful in one evolved clone each. In this case, two repeated P1 transductions yielded no colonies, or colonies that grew very slowly and did not grow within 20 h of dose-response experiments.

### Analysis

#### Estimation of the fold-increase in resistance from the evolution experiments

Final IC_50_ values were estimated from the evolution experiments by averaging the values from the last five time points. Initial IC_50_ values were estimated for each evolving population at the time point when clear inhibition was first observed. Then, each IC_50_ was calculated based on the current growth rate and the corresponding antibiotic concentration using the Hill function as in the experiment. For TMP and NIT, a threshold for the normalized growth rate of 0.55 was used and three consecutive time points from first inhibition were averaged. For MEC, a higher growth-rate threshold of 0.6 was used, since the initial drop in growth rate often remains high, and only this single time point was used. For each gene-deletion strain, all initial IC_50_ estimates from replicate populations were averaged. Both estimations were validated for a subset of gene-deletion strains using direct measurements (Fig. S2).

#### Variability measures

To quantify variability (Fig. 1 and Fig. 2, Fig. S4), we used two different quantities. Resistance Variability (RV) quantifies the variability in the distribution of resistances as a function of time. Pairwise separation (δ) quantifies the relative separation in orders of magnitude between pairs of IC_50_ trajectories over the total duration of the evolution experiments.

<u>Resistance variability measure:</u> RV (labelled *variability* in Fig. 1 and Fig. 2) is defined as the interquartile range over the evolving populations, normalized by the final fold-increase of the median in logarithmic space, i.e., 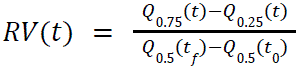, where *Q_i_* are the respective quantiles calculated on the logarithm of the resistance. In other words, this measure quantifies by how many orders of magnitude the resistance increase among the evolving populations is varying in relation to the variation over time.

<u>Pairwise separation δ:</u> Given a pair of IC_50_ time series, *a*(*t*) and *b*(*t*), with *n* time points, we define the pairwise separation in orders of magnitude, δ(*a*, *b*), as 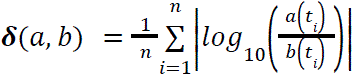, where *t_i_* is the time at dilution *i* and *n* is the total number of dilutions in the experiment (Table 1). Thus, the pairwise separation δ(*a*, *b*) measures by how many orders of magnitude in resistance increase the IC_50_ trajectories *a*(*t*) and *b*(*t*) differ on average.

To avoid artifacts due to noise in the data (mostly due to the technical limitation described in section *Automated high-throughput morbidostat evolution experiments*), we smoothed all individual trajectories (Python *gaussian_filter1d* with FHMW = 4.2 h) before calculating their separation δ.

To plot (Fig. S4) and analyze the distributions of the separations δ of the resistance trajectories of the same ancestral strain, we weighted each distance by the inverse of the total number of distances for that strain. This ensured that each strain contributed equally and that strains with many replicates (such as the reference strain) were not overrepresented.

To determine the significance of the differences in the separation distributions, we used the statistics of the Kolmogorov–Smirnov test. Since the above separations are not uncorrelated, we used 10,000 instances of a permutation null model, in which we randomly assigned time series to the investigated groups (knockout strains, UTI strains) and subgroups (individual strains). Importantly, all results remain unchanged when all time points at which some populations reach the solubility limit are excluded.

#### Analysis of percentage of resistance increase explained by five components

To determine the extent to which the fold increase in resistance can be explained by each of five components (Fig. 2h–j) – initial resistance, initial growth rate, other particularities of the initial genotype, evolutionary stochasticity, and measurement error – we built five linear models that subsequently include each of the five components^8^. We then assessed the increase in the coefficient of determination when fitting each model with respect to the previous model.

Specifically, let *y*_*jkl*_ denote the *l*^th^ measurement of the fold-increase in resistance of population *k* that evolved from ancestor *j*, where *j*∈{1, …, 258} indexes the different ancestors (gene-deletion strains), *k*∈{1, … *K*(*j*)} indexes the replicate populations that evolved from the same gene-deletion strain *j*, and *l*∈{1, …, 5} refers to each of the last five IC_50_ data points used to estimate the final IC_50_ of the populations. The latter serves to capture measurement noise under the assumption that changes of resistance on these time scales are unlikely that late in the experiment. The five nested models are built as follows, where the parameters estimated by the previous model are fixed for each model:

Model 1. Initial resistance of the gene-deletion strain.

*y*_*jkl*_ = α *x*_*jkl*_ + β, where *x*_*jkl*_ = *x*_*j*_ ∀ *k*, *l* is the initial resistance of the gene-deletion strain *j*, and α and β are the model parameters.

Model 2. Effect of initial genotype beyond initial resistance.

Model 2A. Initial resistance and initial growth rate of the gene-deletion strain.

*y*_*jkl*_= α *x*_*j*_ + β + ρ *g*_0*jkl*_, where *g*_0*jkl*_= *g*_0*j*_ ∀ *k*, *l* is the growth rate of the ancestor. This model addresses the question of how much the predictions of model 1 can be improved by correcting the predicted values by a number proportional to each ancestor’s growth rate. Thus, it thus has only one parameter, ρ. Since our selection of ancestral gene-deletion strains already excludes those with strong growth defects compared to the reference strain, this model primarily acts as a sanity check to identify undesired effects on resistance increases that are due to differences in initial growth rates.

Model 2B. Initial resistance, initial growth rate, and initial genotype of the gene-deletion strain. 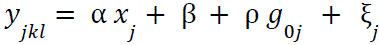. This model has one parameter ξ*_j_* per gene-deletion strain, that accounts for any other properties of the initial genotype (other than the already considered resistance and growth rate, which were already considered in the previous models) that could contribute to an increase in resistance.

Model 3. Initial resistance, initial growth-rate, initial genotype of the gene-deletion strain, and stochasticity of evolution. 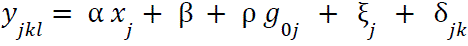. This model has a parameter δ_*jk*_ for each population *k* descended from ancestor *j*, which accounts for differences in fold-increase within populations evolved from the same ancestor. Thus, it represents stochasticity of evolution rather than the idiosyncrasies of ancestors.

Model 4. Initial resistance, initial growth rate, initial genotype of the gene-deletion strain, stochasticity of evolution, and measurement error. 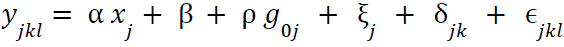. Finally, this model adds the necessary constant ɛ*_*jkl*_* for each data point to be fit exactly. This assumes that all remaining differences between the data and the values predicted by model 3 are due to measurement errors.

All models were fit using the Python function `scipy.linalg.lstsq`, except for Model 4, which has 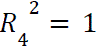 by definition. The coefficients of determination 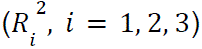 were computed to build the pie charts in Fig. 2h–j.

This analysis only included gene-deletion strains in which at least one population survived for all antibiotics; therefore, all three populations descended from *ΔrelA*, *ΔtolQ,* and *ΔtolB* were excluded.

#### Statistical analysis of IC_50_-versus-time trajectories (for Fig. 4)

To determine which gene-deletion strains had IC_50_ time series that differed from those of the reference strain, we defined a score statistic based on the area between the median trajectory of the reference strain populations and that of every other population. Trajectories were compared in logarithmic space. Then, we performed a Mann–Whitney *U* test on this statistic (Python function *scipy.stats.mannwhitneyu*) with a control of the false-discovery rate (Python function *scipy.stats.false_discovery_control*). To avoid artifacts due to noise in the data, we used the smoothed data for the individual trajectories (same as for variability measure above: Python *gaussian_filter1d* with FHMW = 4.2 h).

Similar results were obtained using several window sizes and smoothing functions (in particular the rolling mean pandas.rolling). In some cases, *ΔfumA* was also significant for TMP (with the current window size *fumA*, *p*=0.05). The same smoothing function (*gaussian_filter1d*) was used in Fig. 4b–c for clearer visualization.

#### Selection of samples for sequencing

For each of the three evolution experiments NIT_evo, TMP_evo and MEC_evo, a priority list was established using a simplified version of the score defined in the previous section. The list was curated manually for each experiment, and all three evolved populations from the first about thirty gene deletions listed were sent for sequencing. Thus, our selection of sequenced populations is biased towards strains in which at least one replicate population has an IC_50_ time series that differs strongly from that of the reference populations. The ancestral strain was also sequenced for each selected gene deletion to distinguish mutations that occurred during the evolution experiments from those that were already present before evolution.

The selected populations were re-grown overnight (for up to 16 h) from the frozen plates in a minimum concentration of the corresponding antibiotic (NIT 3 µg/ml, MEC 0.5 µg/ml, TMP 1.5 µg/ml). The frozen plates correspond to one of the time points at (or close to) the end of the experiment. In most cases, the selected time point is included in the five averaged time points considered in Fig. 2. However, for three samples, growth from those plates failed, so the plate from the previous day was used (Table S1).

#### Analysis of sequencing data

All selected samples (Table S1) were sent for DNA isolation and Illumina NGS whole-genome sequencing (INVIEW resequencing at Eurofins Genomics GmbH – 150 bp paired-end reads, NovaSeq 6000 S4 PE150 XP, unique dual indexing; output: approximately 5 million read pairs; guaranteed minimum 100× average coverage.). Raw fasta files were first pre-processed using Trimmomatic-0.39 and the resulting trimmed files were analyzed using Breseq v0.39.0^79^. Reads were aligned to the Keio parent sequence (Accession: CP009273) using Bowtie2 version 2.5.4. Before running Breseq, IS element annotations were added to the reference genome using ISEscan^80^ to improve junction predictions. We used Breseq in polymorphism mode as the samples were regrown from frozen stocks of evolving populations rather than selecting individual clones (see previous section).

Breseq output files were further inspected and curated using gdtools and custom Python scripts (available upon request). In particular, we systematically analyzed the coverage signal of all sequenced samples to identify amplifications and missing coverage, which are often missed by Breseq. To rule out contamination, we systematically checked whether the expected gene deletion in the genome exhibited missing coverage. Three samples did not pass this check and were excluded from further analysis (Table S3). To validate the amplifications identified, we examined the tables of unassigned junction evidence from Breseq, considering only those amplifications for which at least one corresponding junction was also present. Our curation process may miss amplifications in repetitive regions, but using a more permissive criterion would also identify spurious amplifications. For the examples highlighted in the main text, we manually inspected the coverage signal for any additional amplifications or deletions that were not identified by our systematic analysis. This was particularly important for two *ΔtolC* populations evolved in TMP (experiment TMP_evo), which have a large amplification including the *folA* gene (Fig. S7).

For all analyzed evolved samples, we removed the mutations already present in the ancestors (Table S1) using *gdtools SUBTRACT*. Although Breseq was run in polymorphism mode, we only considered high-frequency mutations (i.e., mutations present in at least 80% of the population) for all sequenced samples. This is because we were interested in mutations in resistant clones that should have mostly taken over the population under the strong selection in our evolution experiments. Intergenic mutations that do not hit gene promoters and synonymous mutations were excluded from further analysis (7.2% of mutations, Fig. S6).

#### Identification of samples with mutator phenotypes

For each experiment, we identified samples as *mutators* if they accumulated more mutations than expected under a null model using a permutation test: More precisely, we created *M* = 10⁵ instances (surrogates) from a computational null model in which we randomly distributed the total number of mutations across all samples. We then determined the number *n* of surrogates whose maximum number of mutations was at least the number of mutations of the most-hit actual sample and considered this sample a mutator if 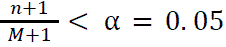 (Table S2). Our null model here entails that the total number of mutations found reflects the mutation rate in the absence of mutators. If we identified a mutator this way, we can retrospectively say that we overestimated the mutation rate in the null model. To avoid false negatives due to this, we therefore repeated the procedure with all previously identified mutators removed from the dataset until it didn’t find any more mutators.

For the evolution experiment with NIT, two of the *ΔuvrD* populations were not classified as mutators by our algorithm (not significant), probably because they did not accumulate a sufficient number of mutations within the duration of the experiment. These samples were manually added to the mutator list, since *ΔuvrD* was present in mutator lists of TMP and MEC, and *uvrD* is known to be involved in mismatch repair^49^.

#### Identification of multihit genes

Through a similar procedure, we identified genes that are hit by mutations in more samples than expected by chance. We counted the number of hits in a specific gene as the number of samples with at least one mutation in that gene (i.e., multiple mutations in the same gene within one sample were only counted once). To simulate the null model, we distributed the total number of hits in each sample over all genes in the genome, with probabilities proportional to their sizes (we added 150 bp to each gene to account for promoter regions, as done by *Breseq*). Then, we identified multihit genes analogously to mutators in the previous section (again with *M* = 10^5^ simulations of the null model and α = 0. 05). These genes are mutated more often than expected by chance and are therefore likely to contribute to the evolved resistance to each antibiotic (Fig. 3; Fig. S8).

#### Parallelism and convergence indices

We use the convergence and parallelism indexes defined in^8^ to quantify whether the degree of repeatability observed in our parallelly evolving populations is greater than expected by chance:

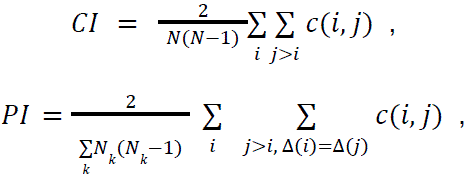

Here, *c*(*i*, *j*) is the number of genes mutated in both evolved populations *i* and *j*, Δ(*i*) is the ancestral gene-deletion strain from where population *i* evolved, *N* is the total number of populations and *N_k_* the number of replicate populations that evolved from ancestor *k*. Thus, *CI* quantifies the degree of convergent evolution between two randomly picked populations (how similar their fixed mutations are at the gene level), while *PI* quantifies the degree of convergent evolution between two populations that evolved from the same ancestral gene-deletion strain.

For our morbidostat evolution experiments with strong selection, the observed values of *CI* are significant when compared with a null distribution where mutations are randomly distributed through all genes in the genome with a probability proportional to their size (*p*=10^−4^ for all antibiotics). For *PI*, we simulated the null model by randomly shuffling the information on the initial gene deletion in the *c*(*i*, *j*) matrix, but keeping its structure (permutation test). The observed *PI* value was statistically significant for all antibiotics (Fig. 5a; TMP *p*=0.048, MEC *p*=10^-4^, NIT *p*=0.0068).

#### Statistical analysis of inhibitors effect on resistance increase

We asked whether small-molecule inhibitors that inhibit the gene-functions identified as modifiers of resistance (Fig. 4) could have similar effects on resistance evolution in genetic backgrounds containing no genetic modification of these genes. We thus performed experiments with six conditions and several strains (Fig. S13), and addressed the question on whether there are any statistically significant trends of the inhibitors to reduce resistance increase on the relevant strains. For each condition, we performed a one-sided *t*-test to obtain a *p* value for each UTI strain and the reference strain, and then combined those *p* values using Mudholkar’s and George’s method^81^ to assess whether the observed global trend was significant. We further corrected the combined *p* values for all six conditions using Benjamini’s and Hochberg’s false-discovery control. For the analysis at the level of individual strains (red dots in Fig. S13), we performed a two-sided *t*-test and again controlled the false-discovery rate with Benjamini’s and Hochberg’s adjustment. When comparing two different experiments, the fold increase at the final time point of the shorter experiment was used. Python functions used are all from `scipy.stats`; for the one-sided *t*-test on individual strains `ttest_ind,` we used *equal_var=False*.

## Acknowledgments

We thank the whole Bollenbach group, M. Lukačišinová, J. Krug, and M. Lässig for fruitful discussions; M. Lässig for critical reading of the manuscript; L. Goretzky for technical support; the High Performance Computing (HPC) team at University of Cologne for support with the cluster; the HighRes Biosolutions team for support with the robotic system; J. Barrick for support with Breseq and A. Baurmann for the schemes in this manuscript. G.P. thanks N. Freitas, A. Baurmann and her friends in Cologne for support of all other kinds. This work was supported by German Research Foundation (DFG) Collaborative Research Centre (SFB) 1310.

## Author contributions

Conceptualization: T.B., G.P., and T.F. Investigation: G.P. designed and developed the protocols and scripts for the implementation of the automated high-throughput morbidostat in the robotic system, and performed the evolution experiments *NIT_evo* and *TMP_evo*, G.P. and T.F. designed and performed the evolution experiments *MEC_evo* and *inhibitors_evo* and all other experiments, B.F. constructed the gene deletions in the evolved backgrounds (Fig. 5g–h and S12). Formal Analysis: G.P. and T.F. analyzed the data; statistical analysis were performed jointly by G.P., T.F. and G.A.. Writing – original draft: T.B. and G.P.. Writing – review and editing: T.B., G.P., T.F., and G.A. Supervision: T.B. Funding acquisition: T.B.

## Supplementary figures

**Fig. S1:**
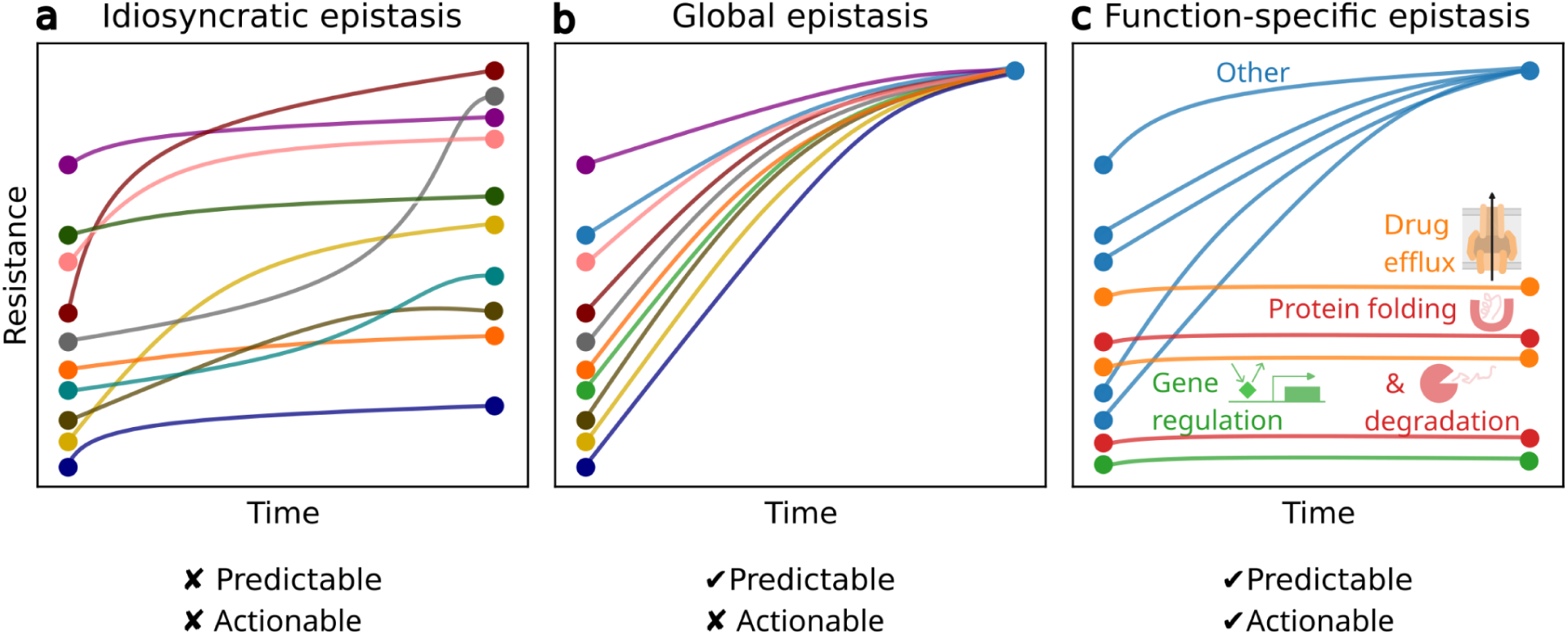
Schematic comparison of the different types of epistasis mentioned in this study. Unlike global epistasis patterns, where evolutionary increases in resistance can be predicted based solely on initial resistance values, or idiosyncratic epistasis, where evolutionary outcomes are unpredictable, function-specific epistasis enables partial predictions of resistance evolution based on the functions of disrupted genes. The identified functions may inform targeted actions to improve drug treatments using small-molecule inhibitors.

**Fig. S2:**
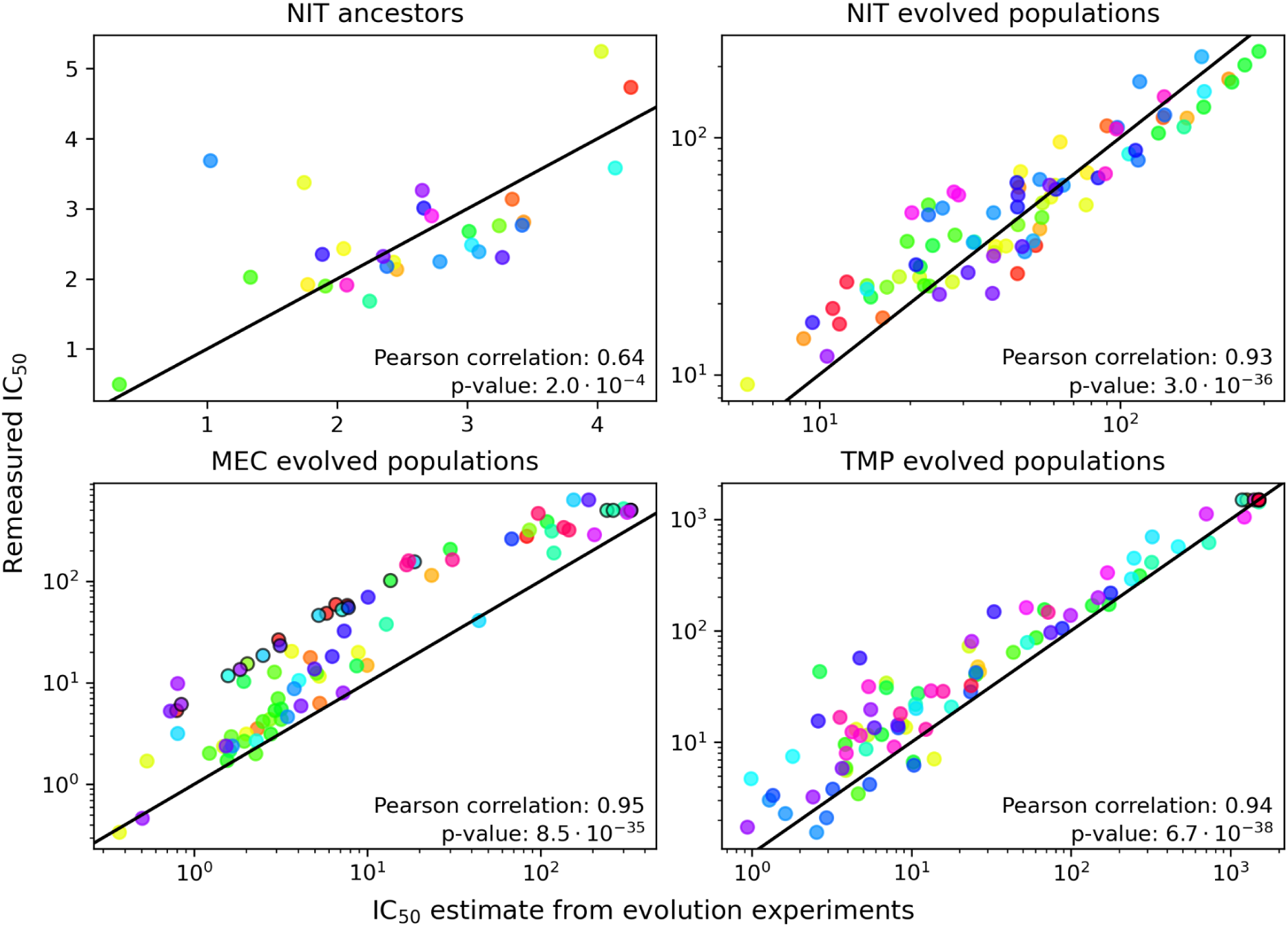
Comparison of IC_50_ values estimated from the evolution experiment with the values remeasured in dose-response curves after the experiment. The data are from experiment NIT_evo (Table 1). Due to the smaller data range, the IC_50_ values of the ancestors are shown on a linear scale. The black lines are the identity. Colors represent different gene-deletion strains. Circles with black edges show the maximum concentration used in the gradient for the dose-response curve measurement and represent a lower limit for the IC₅₀. These cases were excluded from the correlation analysis.

**Fig. S3:**
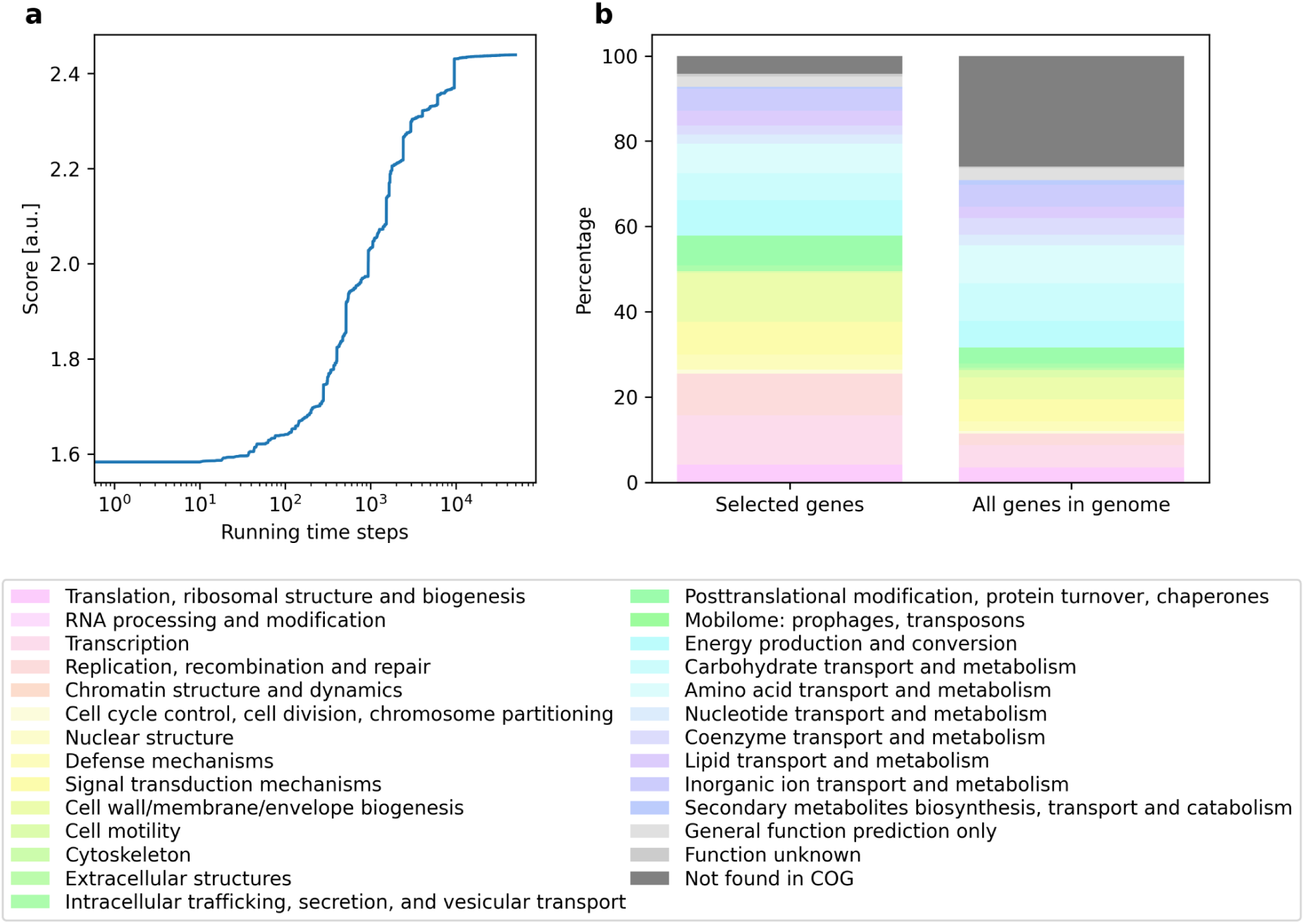
Functional annotation of selected gene-deletion strains. **(a)** Time evolution of the recursively increasing score based on the standard deviation of initial resistance levels for the three antibiotics considered in this study (Methods, section *Selection of gene-deletion strains for evolution experiments*). **(b)** Distribution of COG (Clusters of Orthologous Genes) for the selected genes, compared to the distribution of all genes in the genome.

**Fig. S4:**
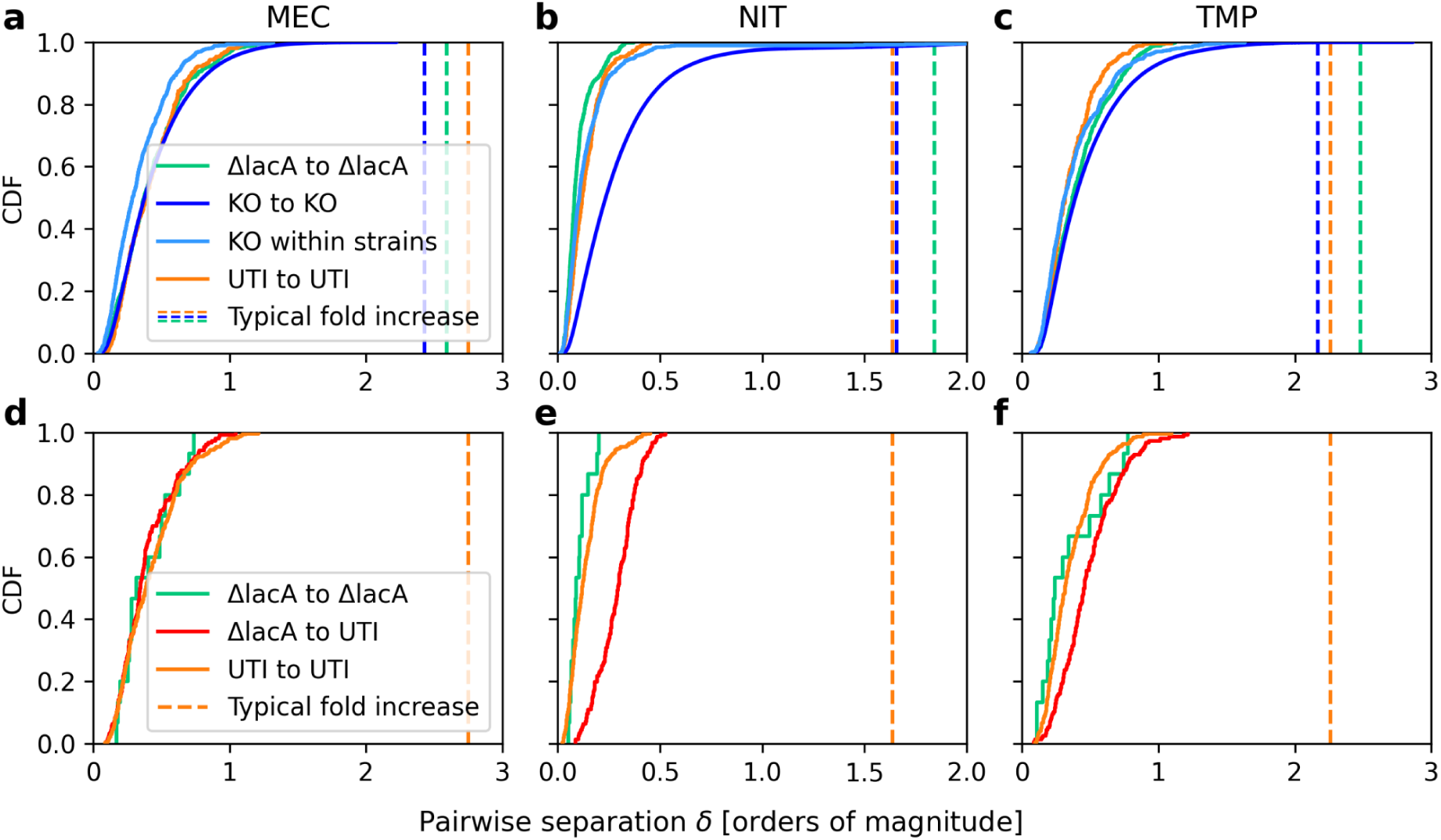
Distributions of the pairwise separation between resistance trajectories. (**a–c**) Pairwise separations δ between IC_50_ time series of gene-deletion strains (KO), either for all pairs (blue) or grouped by trajectories within populations that evolved from the same ancestral gene-deletion strain (light blue, weighted to give each strain an equal total weight), reference strain (*ΔlacA*; green) and clinical isolates (UTI; orange), respectively (Methods). Note that the sample size for the UTIs is too small for a meaningful within-strain distribution. The vertical lines show the median fold increase for the relevant strains (UTI isolates or gene-deletion strains, respectively) over the total duration of the experiment. Almost all values are far below these fold increases, showing that the phenotypic variability is generally much smaller than the typical resistance increase during evolution. Permutation tests show that the distributions for gene-deletion strains and UTI isolates significantly differ for NIT (*p*<0.001) but not for MEC and TMP (*p*=0.68 and *p*=0.051, respectively; Methods). For each antibiotic, the distributions of separations δ between all gene-deletion strains strains (blue) differ significantly (*p*<0.001, Methods) from those within gene-deletion strains (light blue). **(d–f)** Pairwise separation between UTI and reference *ΔlacA* strains (red) and between trajectories within these groups (UTI, orange; *ΔlacA*, green). For MEC, the *ΔlacA* trajectories are as close to the UTI trajectories as the UTI trajectories are to each other (permutation test yielded indistinguishable red and orange distributions, *p*=0.33). For NIT and TMP, trajectories of UTI strains and the reference strain are closer within these groups (orange and green) than across the two groups (permutation tests orange to red, *p*<0.01). However, the difference is much smaller than the typical increase over time (vertical line).

**Fig. S5:**
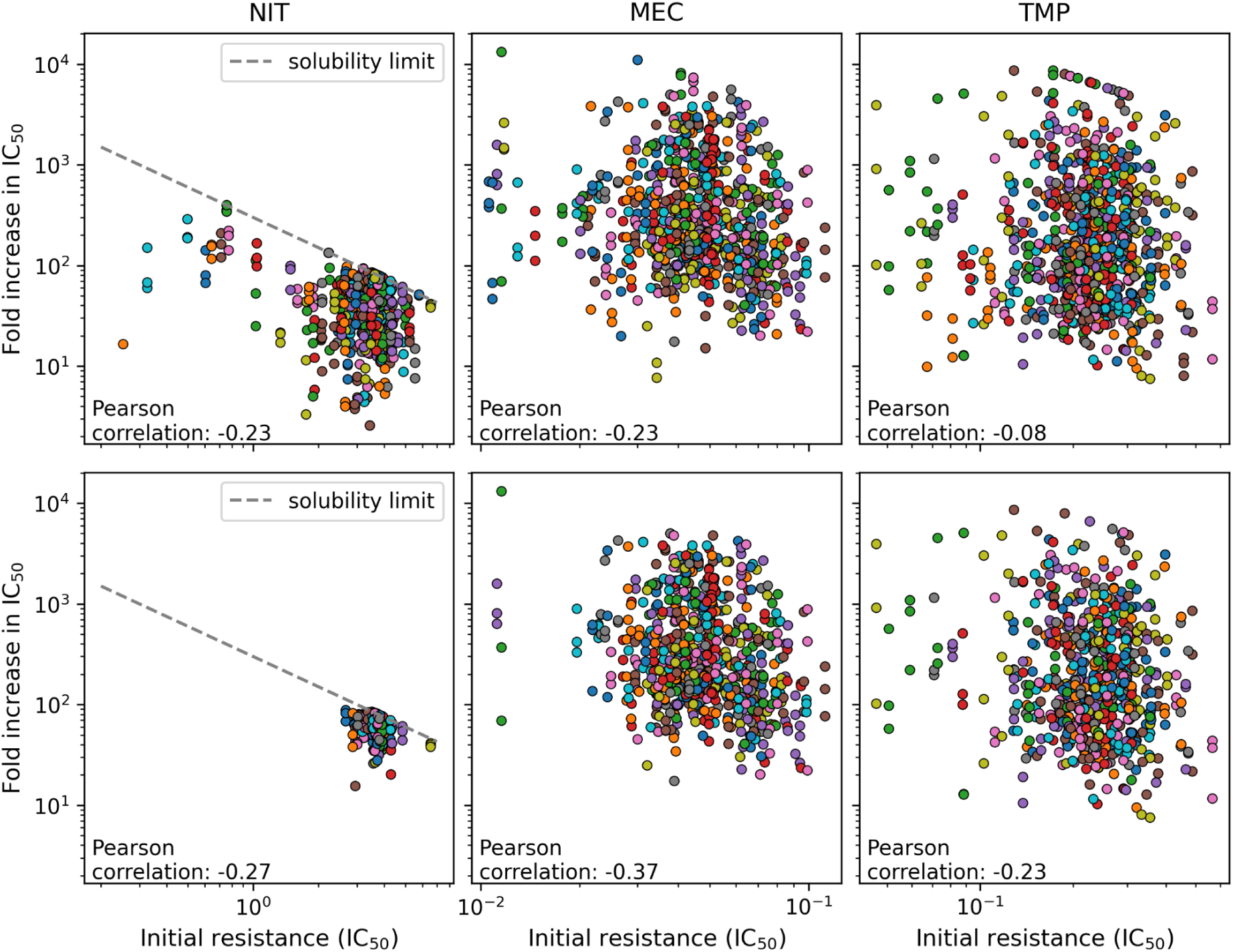
Diminishing returns epistasis does not adequately explain the increase in resistance. Upper panels: Fold-increase in resistance versus initial resistance (IC_50_) for all populations (i.e., 259 gene-deletion strains including the reference strain) in the three evolution experiments NIT_evo, MEC_evo and TMP_evo (Table 1). **Lower panels:** As upper panels, but excluding strains that significantly alter the evolution of resistance (significant gene deletions in Fig. 4 for each antibiotic). Pearson correlation was calculated based on the means of replicate populations from the same gene deletion to avoid pseudoreplication.

**Fig. S6:**
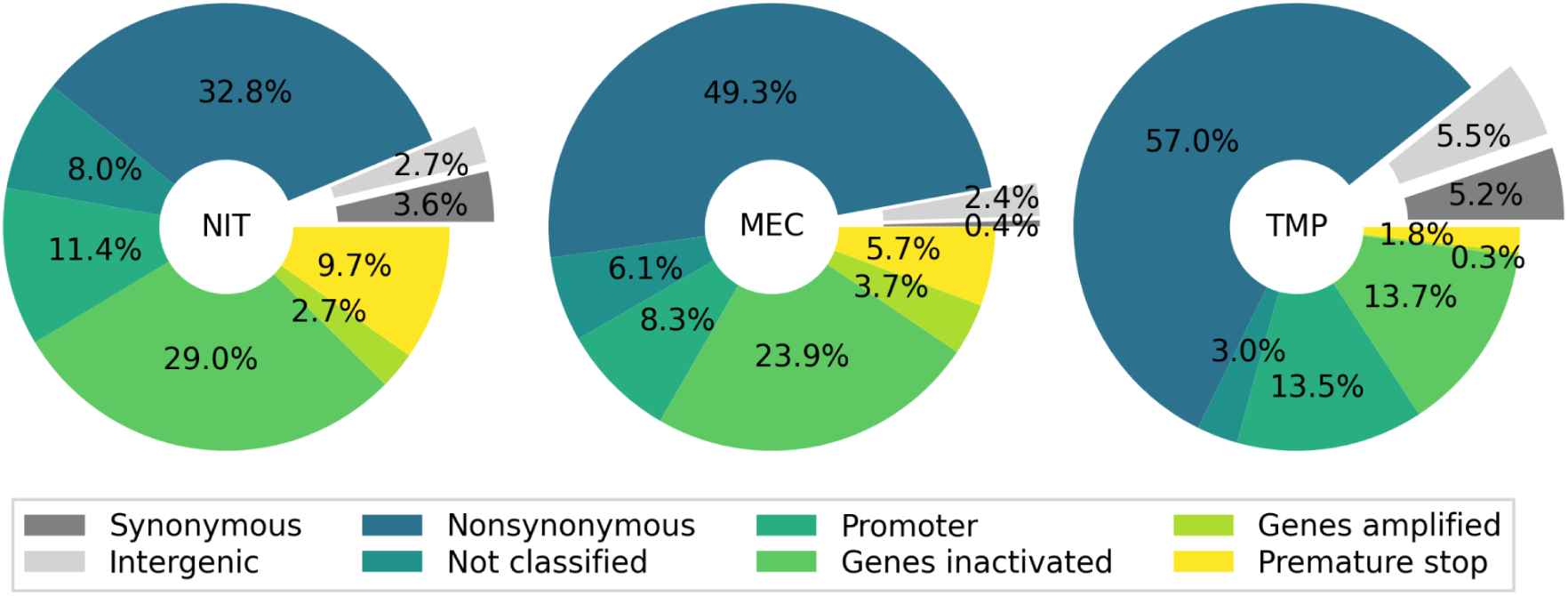
Fixed mutations in sequenced evolved samples. Intergenic and synonymous mutations were excluded from further analysis. See *Analysis of sequencing data* in Methods for details.

**Fig. S7:**
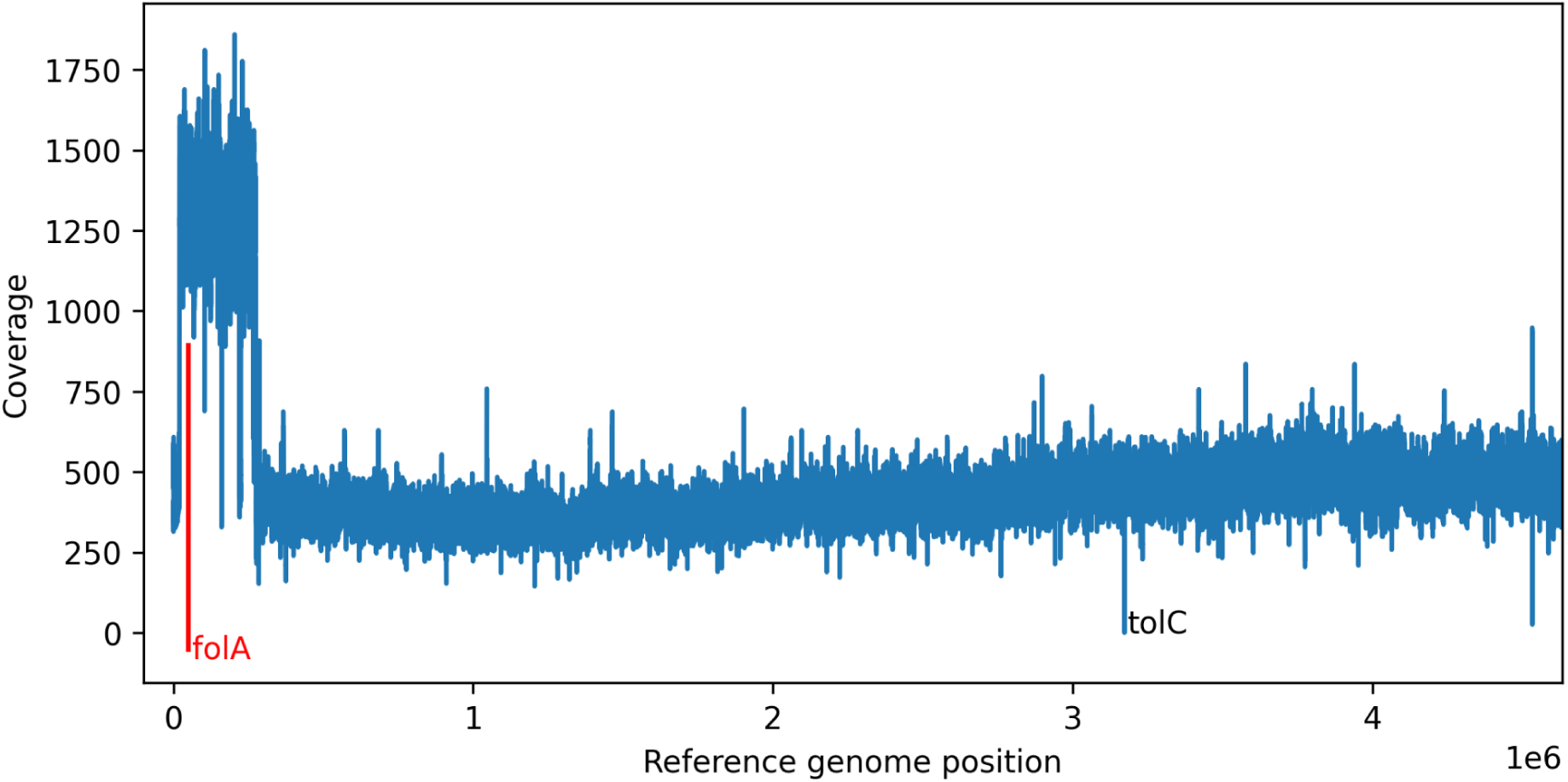
Example of sequencing coverage analysis. Data are from a *ΔtolC* strain that evolved in the presence of TMP (sample 11642_end_F9_tolC, Table S1). A large amplification includes the *folA* gene, which encodes the target of TMP. See *Analysis of sequencing data* in Methods for details.

**Fig. S8:**
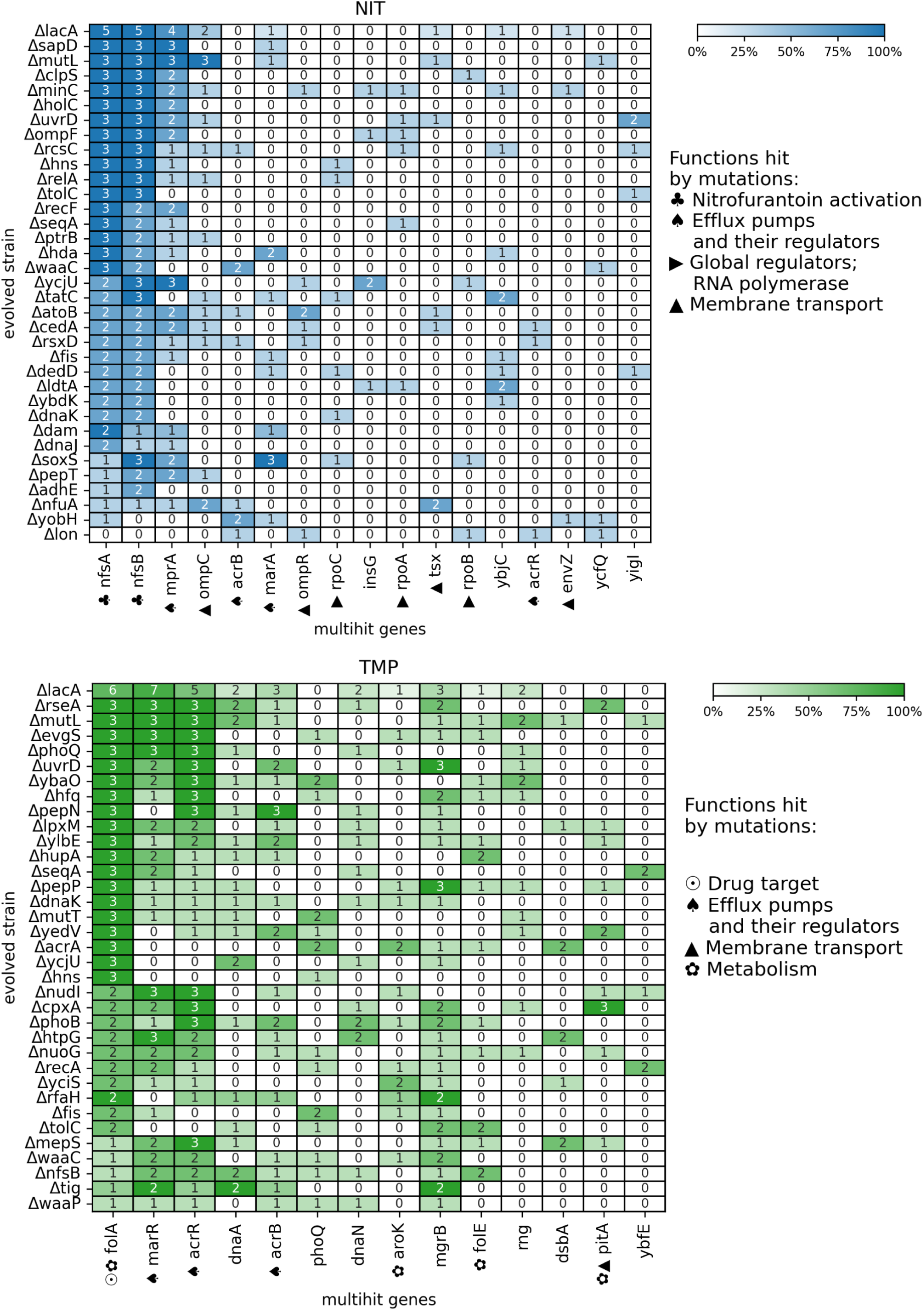
Overview of the mutations in multihit genes fixed in all sequenced evolved populations. As Fig. 3b, but showing the data for NIT and TMP. See *Analysis of sequencing data* in Methods for details.

**Fig. S9:**
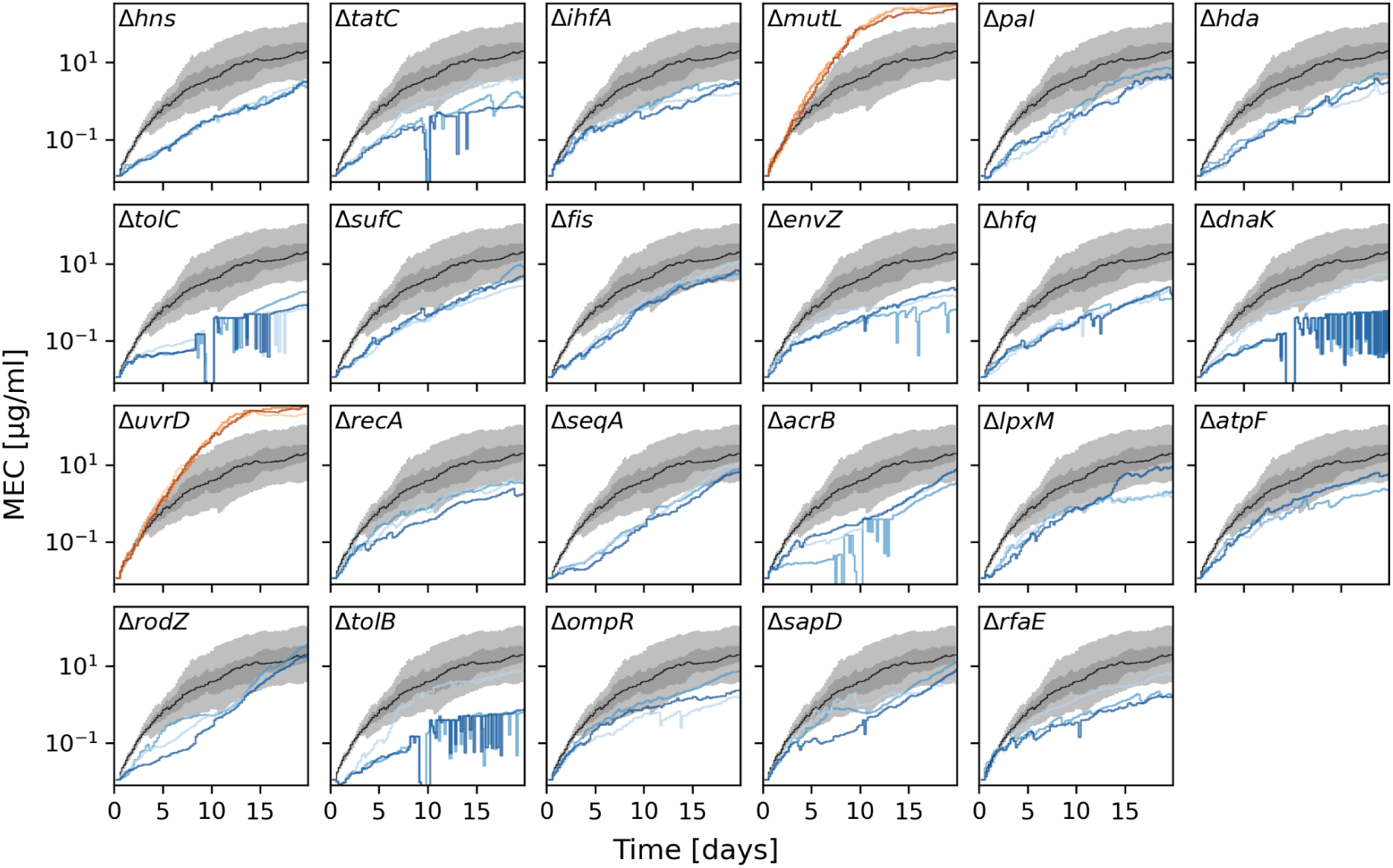
Gene deletions affecting resistance evolution to MEC. Estimated IC_50_ over time from the evolution experiment MEC_evo (Table 1). The dark and light gray shaded areas show the 50% and 90% ranges of IC_50_ values of the reference strain, respectively, and the black line indicates the median. The IC_50_ trajectories of specific gene deletions with significantly increased or decreased resistance evolution are shown in orange and blue, respectively. Different shades indicate three replicate populations, except for the gene-deletion strains listed in Table S4, for which one or two populations died during the experiment.

**Fig. S10:**
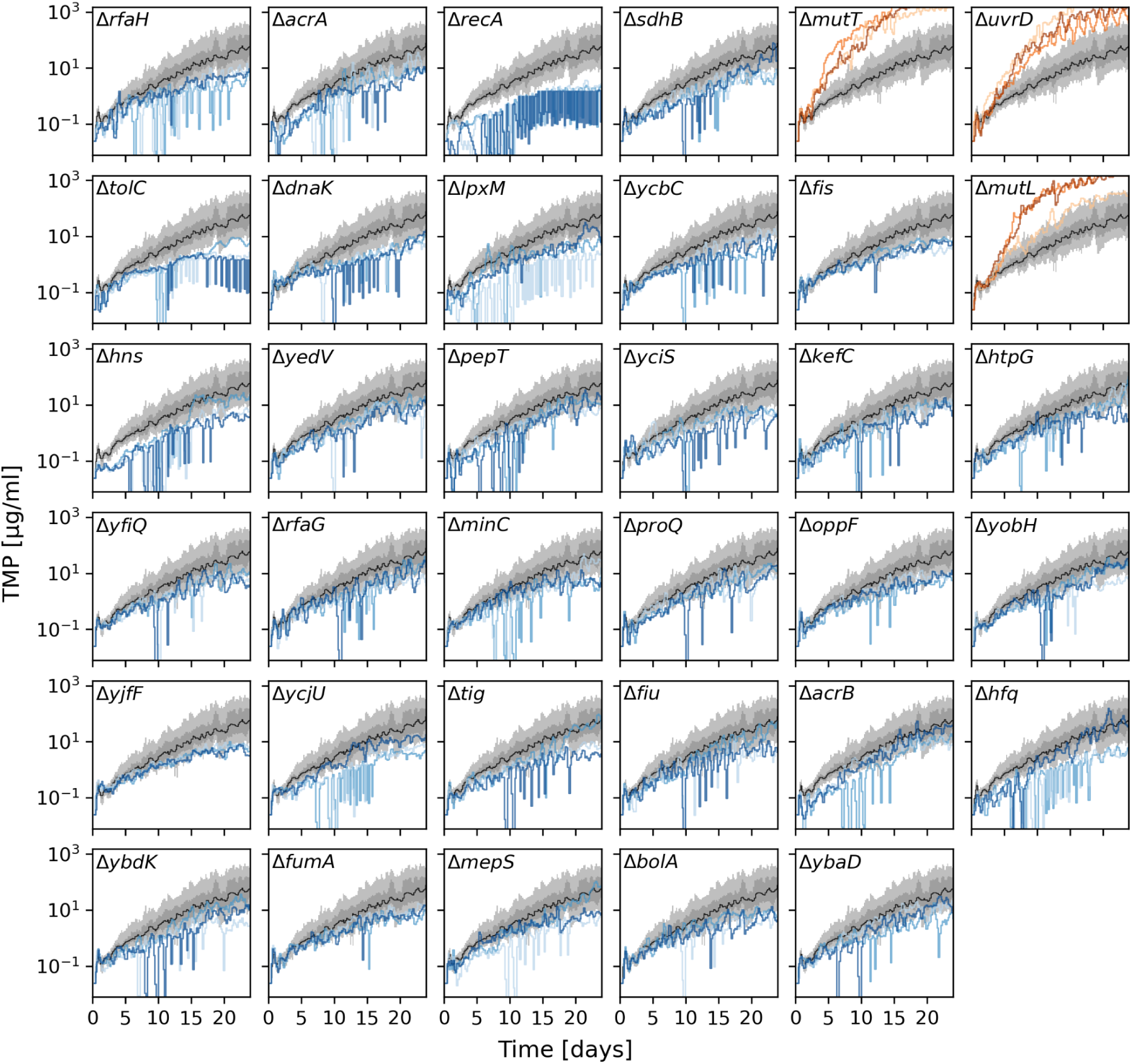
Gene deletions affecting resistance evolution to TMP. As Fig. S9, but showing data from the evolution experiment TMP_evo (Table 1).

**Fig. S11:**
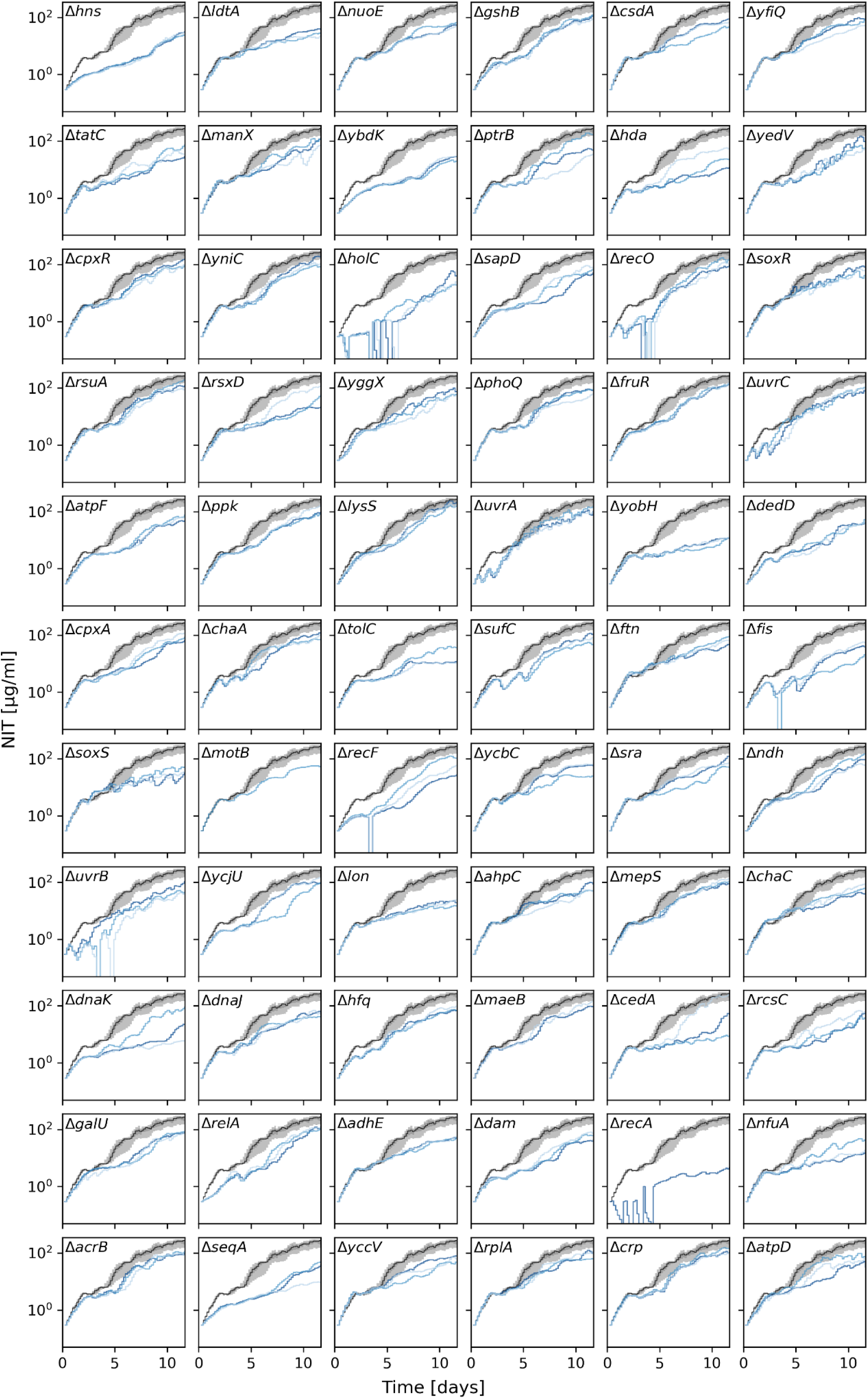
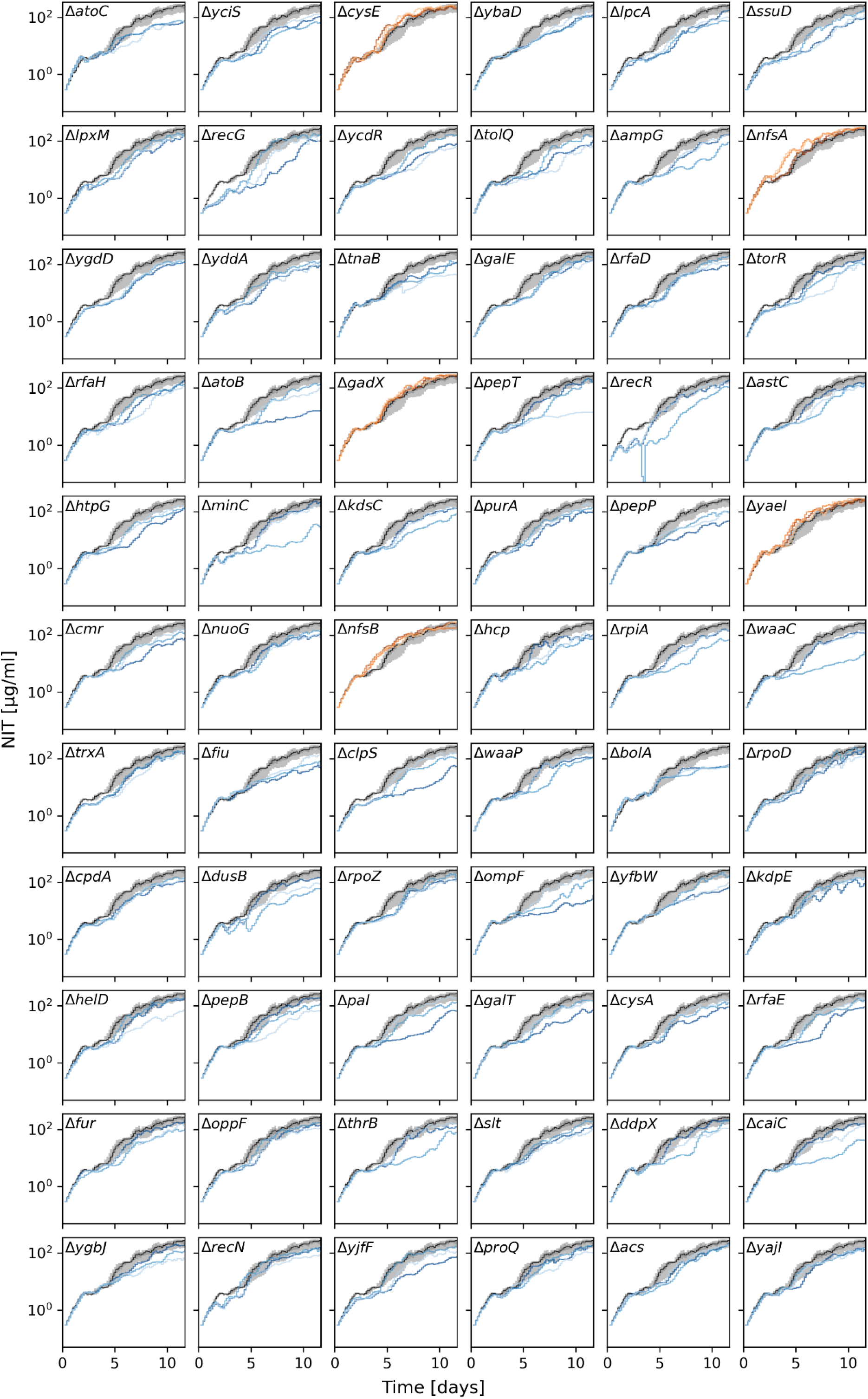
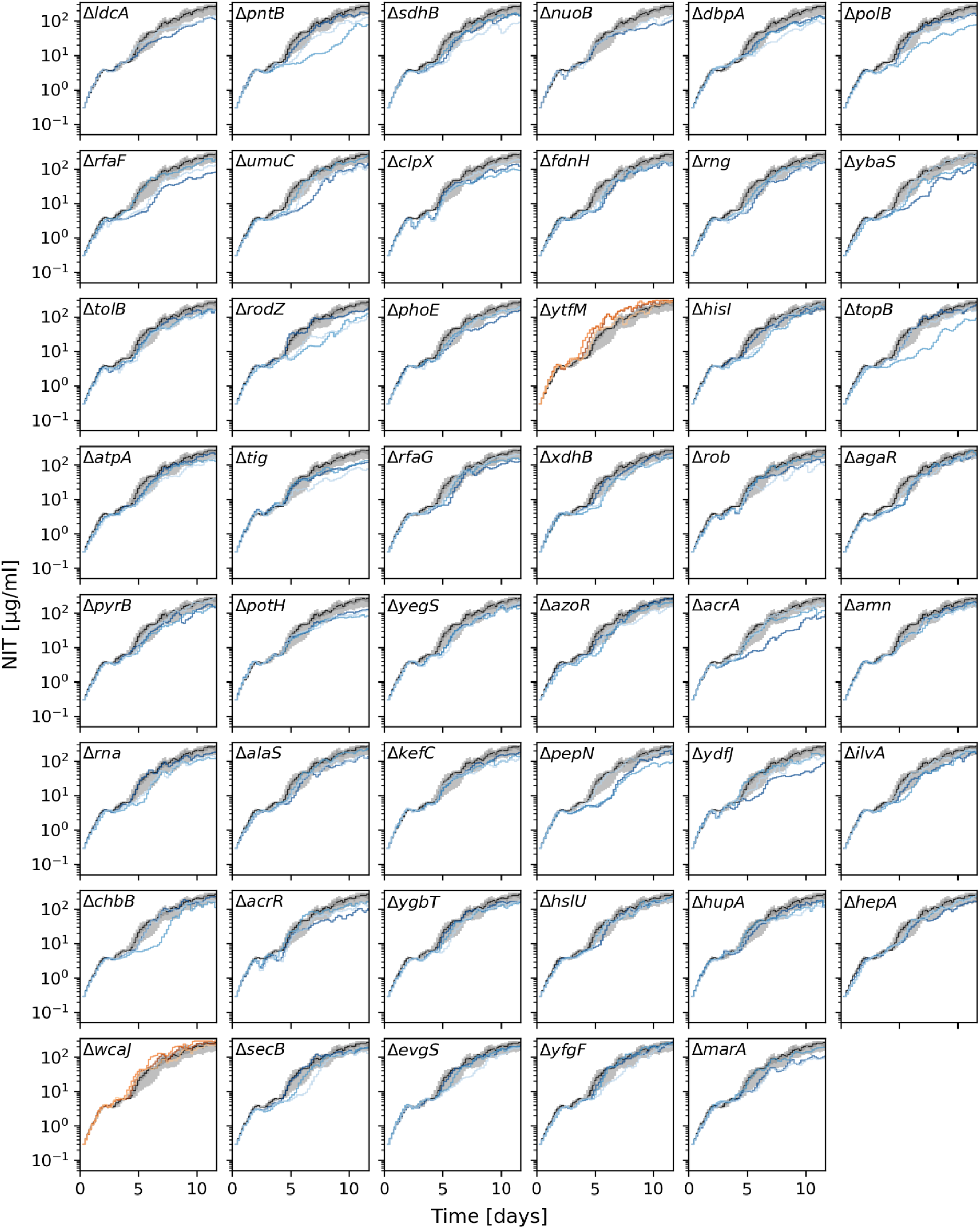
Gene deletions affecting resistance evolution to NIT. As Fig. S9, but showing data from the evolution experiment NIT_evo (Table 1).

**Fig. S12:**
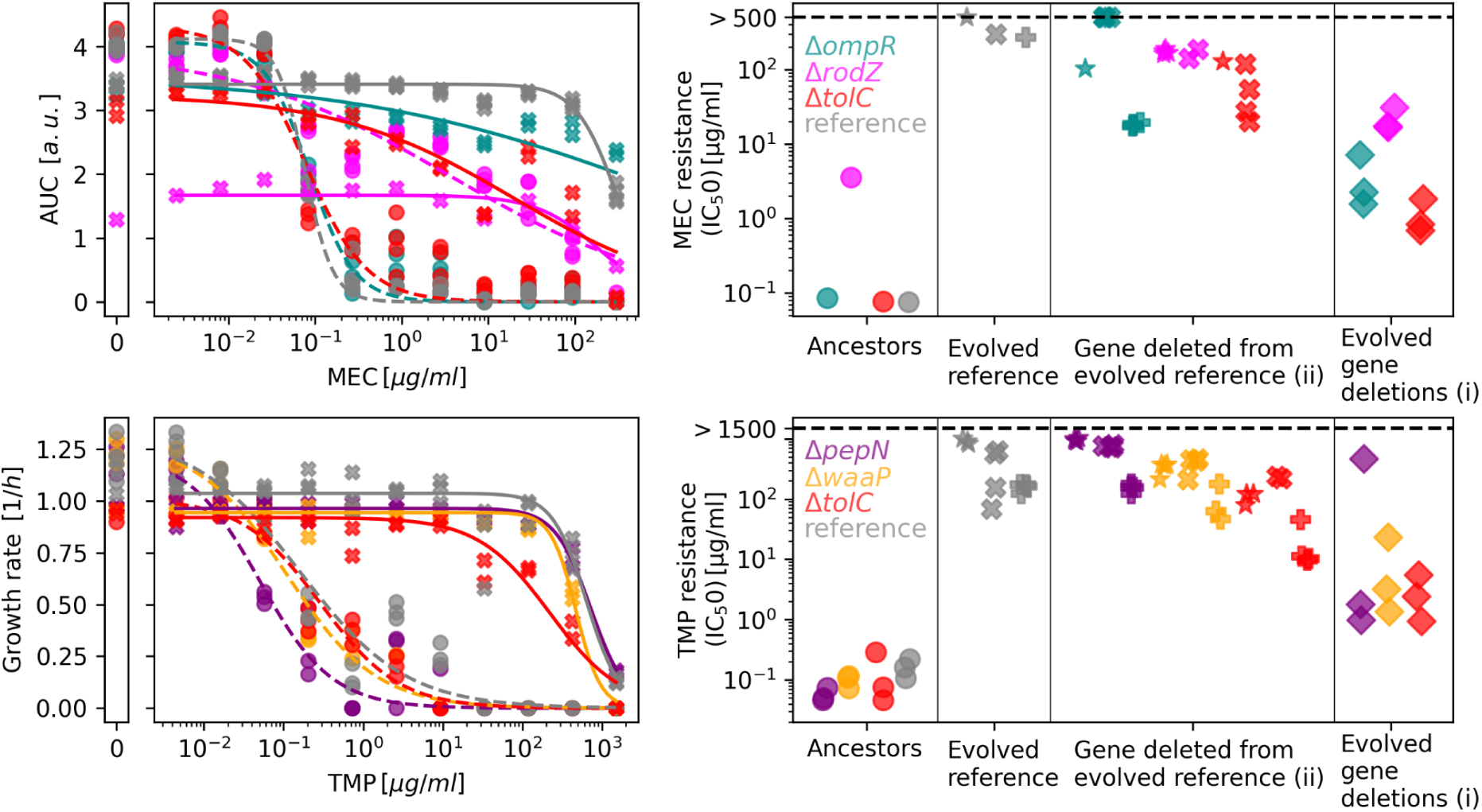
Comparison of (i) the IC₅₀ values for ancestral and evolved strains with (ii) those of the evolved reference strain clones with an additional gene deleted. As Fig. 5g-h, but for MEC and TMP. Symbols and colors in the left panels match those in the right panels.

**Fig. S13:**
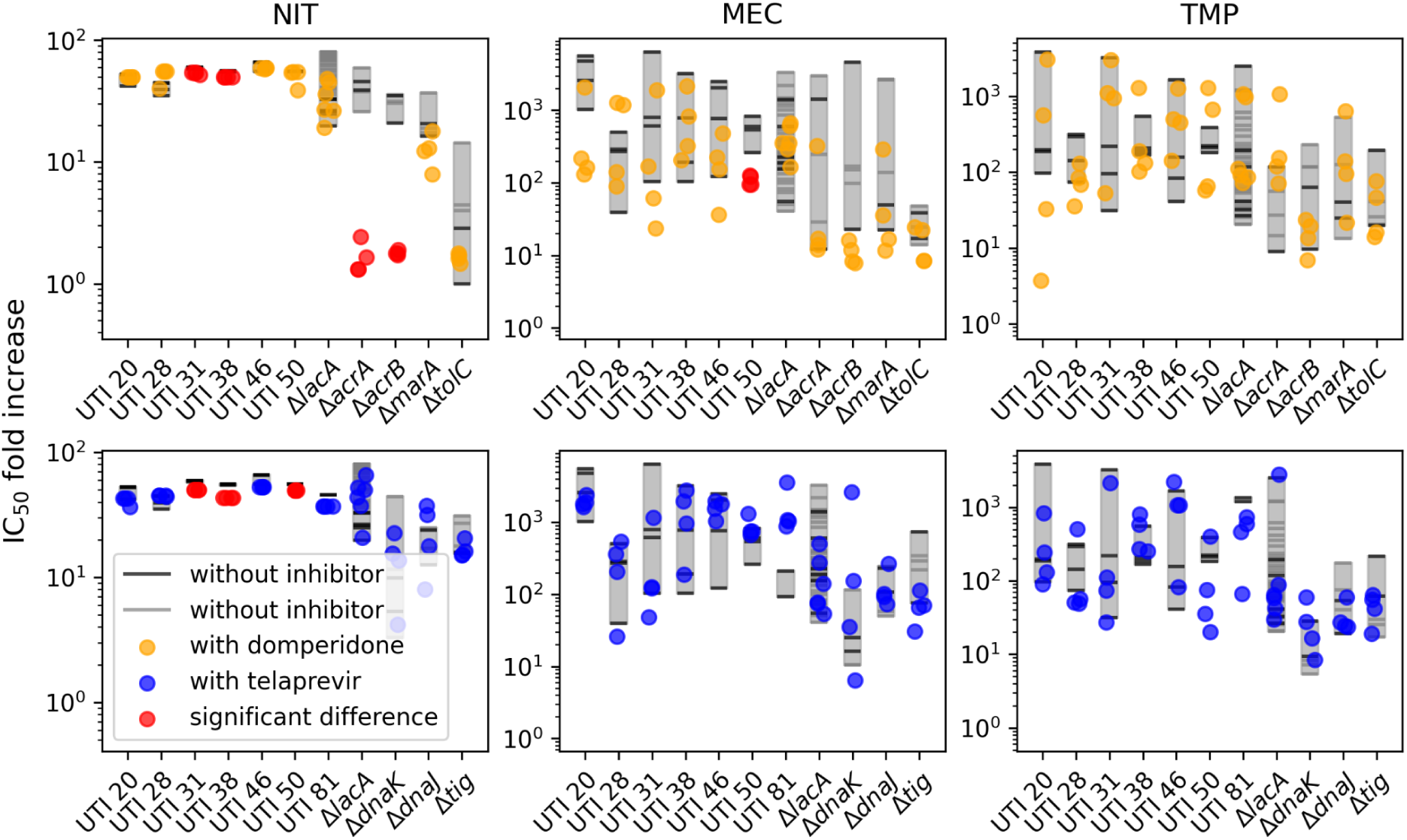
Complete set of comparisons of resistance evolution in the presence and absence of small-molecule inhibitors. The top row and orange points show the fold increase in resistance (IC_50_) in the presence of domperidone (*cf.* Fig. 6d). The bottom row and blue points show the fold increase in resistance in the presence of telaprevir. Gray and black lines show replicates of the reference strain without inhibitor from two different evolution experiments (NIT/TMP/MEC_evo and inhibitors_evo, respectively, in Table 1). Gray rectangles show the range of fold increase for easier comparison. Red dots indicate strains where the inhibitor causes a significant reduction in fold increase (i.e. *p*-values <0.05 in two-sample *t-*tests using Benjamini–Hochberg false-discovery control). Note that in the case of NIT, significant differences except for *ΔacrA* and *ΔacrB* are artifacts because these strains reach the maximum NIT concentration with and without the inhibitor, but the initial IC_50_ changes due to the presence of the inhibitor.

## Supplementary tables

**Table S1:** Sequenced samples and their fixed mutations.

**Table S2:** Samples that spontaneously developed a mutator phenotype during evolution. The last part of the sample labeling indicates the deleted gene.

| Sample | Number of mutations | Experiment |
| --- | --- | --- |
| 17226_19347_D5_clpS | 40 | MEC_evo |
| 17221_19353_B1_acrA | 37 | MEC_evo |
| 17221_19353_E3_seqA | 19 | MEC_evo |
| 17222_19356_E3_seqA | 17 | MEC_evo |
| 17223_19358_E3_seqA | 16 | MEC_evo |
| 17225_19346_C2_pal | 16 | MEC_evo |
| 11646_end_E1_nudI | 299 | TMP_evo |
| 11641_end_F5_phoB | 250 | TMP_evo |
| 11646_end_G5_ylbE | 218 | TMP_evo |
| 11644_end_C5_lacA | 150 | TMP_evo |
| 11645_end_A12_evgS | 149 | TMP_evo |
| 11644_end_F10_pepP | 127 | TMP_evo |
| 11647_end_H7_rseA | 114 | TMP_evo |
| 11649_end_H7_rseA | 93 | TMP_evo |
| 11641_end_G7_ybaO | 80 | TMP_evo |
| 11641_end_H2_hupA | 57 | TMP_evo |
| 11641_end_F9_tolC | 37 | TMP_evo |
| 11643_end_G7_ybaO | 32 | TMP_evo |
| 11645_end_C5_lacA | 26 | TMP_evo |
| 11649_end_C2_lpxM | 25 | TMP_evo |

**Table S3:** Samples with unexpected sequencing coverage in the gene-deletion loci.

| Experiment | Sample | Problem | Excluded? |
| --- | --- | --- | --- |
| NIT_evo | 8962_830_B11_dam | cross-contaminated, another gene-deletion present in missing coverage | yes |
| TMP_evo | 11649_end_B6_rfaH | contamination, no missing coverage at the expected gene position | yes |
| TMP_evo | 11642_end_G4_tig | <10% of median coverage at the expected gene position and expected junction with KAN cassette not present in Breseq output | yes |
| TMP_evo | 11646_end_F8_uvrD | <10% of median coverage at the expected gene position but expected junction with KAN cassette present | no |

**Table S4:**
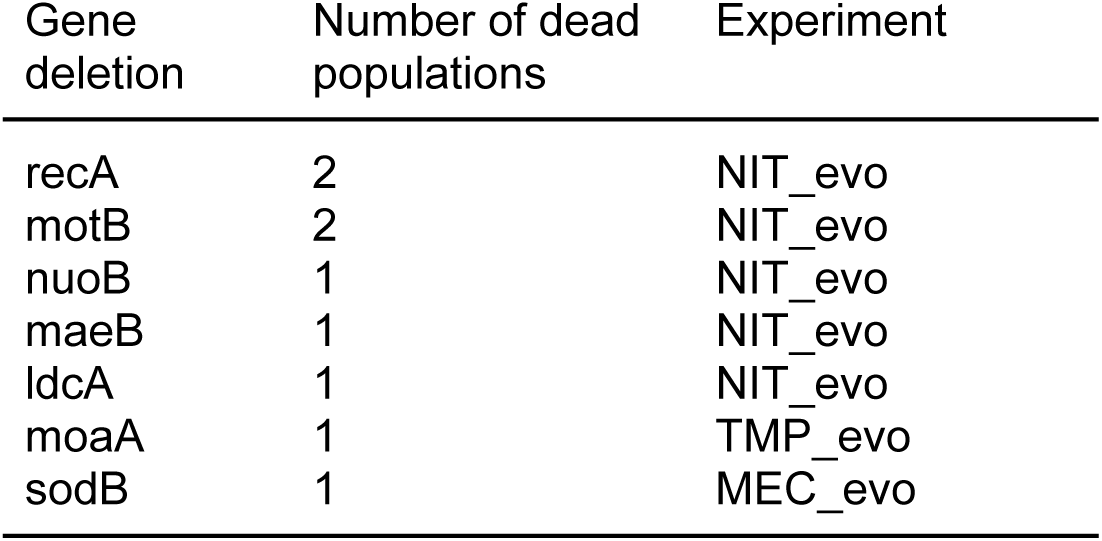
Populations that died during the evolution experiment and were excluded.

**Table S5:**
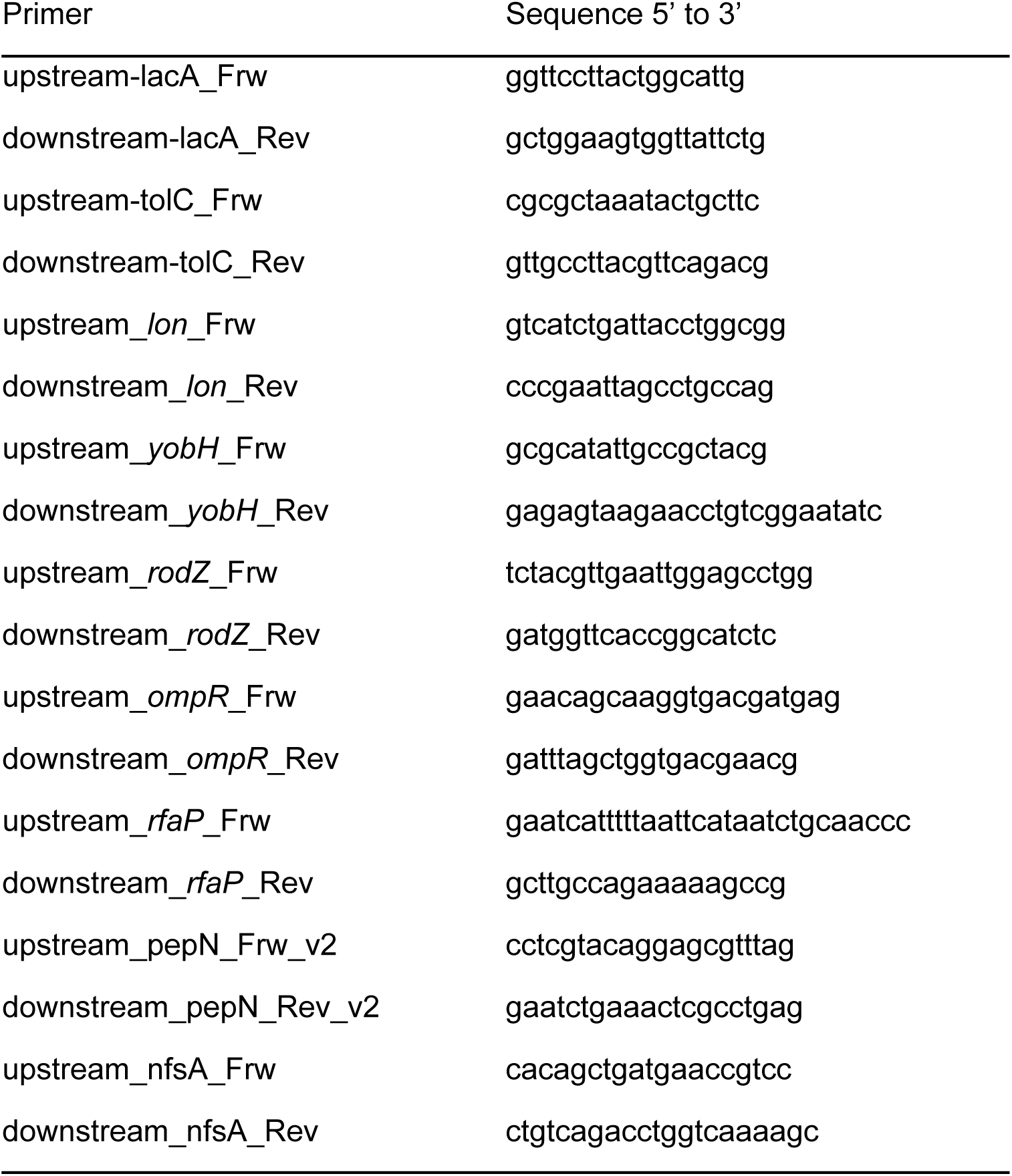
Primers used to generate and verify the deletion of genes of interest in evolved *ΔlacA* clones.

